# Disk-Drive-Like Operations in the Hippocampus

**DOI:** 10.1101/2022.10.05.511000

**Authors:** Wilten Nicola, David Dupret, Claudia Clopath

## Abstract

The rapid computation of re-playable memories within the hippocampus in the form of spike sequences is a near computer-like operation. Information can be encoded once during the initial experience, and replayed numerous times after in a compressed-time representation [1–8]. Theta oscillations, sharp-wave ripples, and attractor dynamics have been posited to collectively play a role in the formation and replay of memories. However, the precise interplay between these dynamical states remains elusive. Here, we show that the memory formation dynamics and operations of the hippocampus are not just computer-like, but map directly onto the dynamics and operations of a disk-drive. We constructed a tripartite spiking neural network model where the hippocampus is explicitly described as a disk drive with a rotating disk, an actuator arm, and a read/write head. In this Neural Disk Drive (NDD) model, hippocampal oscillations map to disk rotations in the rotating disk network while attractor dynamics in the actuator arm network point to “tracks” (spike assemblies) on the disk. The read/write head then writes information onto these tracks, which have temporally-structured spikes. Tracks can be replayed during hippocampal ripples for consolidation. We confirmed the existence of interneuron-ring-sequences, predicted by the rotating disk network, in experimental data. Our results establish the hippocampus as a brain region displaying explicit, computer-like operations. Based on the known interactions between the hippocampus and other brain areas, we anticipate that our results may lead to additional models that revisit the hypothesis that the brain performs explicit, computer-like operations.

## 1 Introduction

The metaphor that the brain operates as a computer has been pervasive in neuroscience since Jon von Neumann’s pioneering work in the 1950’s [9–19]. At the near simultaneous dawn of computer science and electrophysiologial-based, computational neuroscience, Von Neumann postulated that the nearly discrete action potentials fired by neurons were comparable to the digital binary units or bits in the vacuum tubes and transistors of early computers [9]. Such a metaphor, if made into a concrete model, could help reach a comprehensive understanding of, and formulate predictions on, the nature of the computational operations underlying brain functions.

Unfortunately, von Neumann’s efforts were limited by the state of knowledge of the brain at the time. However, the last decades of neuroscientific work have now shed important insights into the cellular substrates and network dynamics at the nexus of brain and behaviour, laying the foundational knowledge about how some neuronal regions specialize and adapt to perform specific operations [20–24]. Notably, the hippocampus of the mammalian brain holds mnemonic information used to inform behaviour [1–8, 20, 25–46]. Likewise, modern computers hold information for further operations using dedicated components: Hard Disk Drives (HDDs) [47].

Accordingly, we test a more direct version of von Neumann’s “brain-as-a-computer” analogy by establishing a theoretical framework where hippocampal operations and dynamics are mapped directly onto those of a computer’s disk drive in the Neural Disk Model of Hippocampal Function. The NDD model is a tri-partite network where each network maps onto the core components of a disk drive: the rotating disk network, the read/write head network, and the actuator arm network. The NDD model also successfully maps theta oscillations and sharp-wave ripples to disk-rotations while attractor dynamics act as the actuator arm, pointing to individual tracks or cell assemblies. The zoo of observed hippocampal replays (forward replay, reverse replay, extended replay, fragmented replay) can interpreted as specific data-accessing events in the NDD model. Finally, we detected the interneuron-ring sequences in multi-unit recordings from rats, as predicted by the rotating disk network.

## 2 Results

### 2.1 Mapping Hippocampal Behaviours to Disk Drive Dynamics

We start by first describing hippocampal dynamics. The spike times of hippocampal pyramidal cells and interneurons are organized on multiple timescales by a collection of network oscillations that are observed as rhythmic fluctuations in the local field potentials (LFPs) and correlate with behavioural states and memory processing stages (Figure 1A). Sequences of said spikes are observed on two time scales: temporally compressed spike sequences during hippocampal sharp-waves ripples (SWRs, Figure 1A-B) and temporally dilated forms of these spike sequences during hippocampal theta oscillations (Figure 1A,C). The SWR LFP event is a 125–250 Hz oscillatory event lasting approximately 100 ms and supporting memory consolidation [1–8, 26, 48–53]. The theta oscillation, on the other hand, is a 5–12 Hz oscillation that dominates hippocampal LFPs, organising temporally structured firing activity of pyramidal cells in support of learning during active exploration (Figure 1C) [20, 25, 27, 44, 54, 55]. During rest/sleep, theta-nested neural patterns are compressed and replayed in sharp wave/ripples (SWRs) [1–8, 56–62]. The relation between compressed sharp-wave sequences and theta sequences may occur through an oscillatory-interference mechanism where one oscillator controls spike times during SWRs, and a second oscillator dilates SWR-sequences into theta sequences by creating an interference pattern that dilates the sequential content of the carrier waves into the envelope phases (Figure 1C) [63]. This postulated interference mechanism provides a mechanistic explanation for hippocampal phase precession [20,44,64–72], and explicitly links theta sequences during a single cycle of the theta oscillation to compressed spike sequences during a SWR [63]. Indeed, spike sequences during a cycle of the theta oscillation are also a compressed representation of the firing fields of individual cells, and are known to have comparable compression rates to replay sequences in SWRs [48]. Hebbian plasticity allows for one-shot learning of new sequences by using existing theta-sequences as a compressible temporal backbone (Figure 1D). Different populations of neurons within a theta sequence can be selected by biasing currents to store potential information (Figure 1E) allowing for discrete memories to be stored in different populations of pyramidal neurons [73].

**Figure 1:**
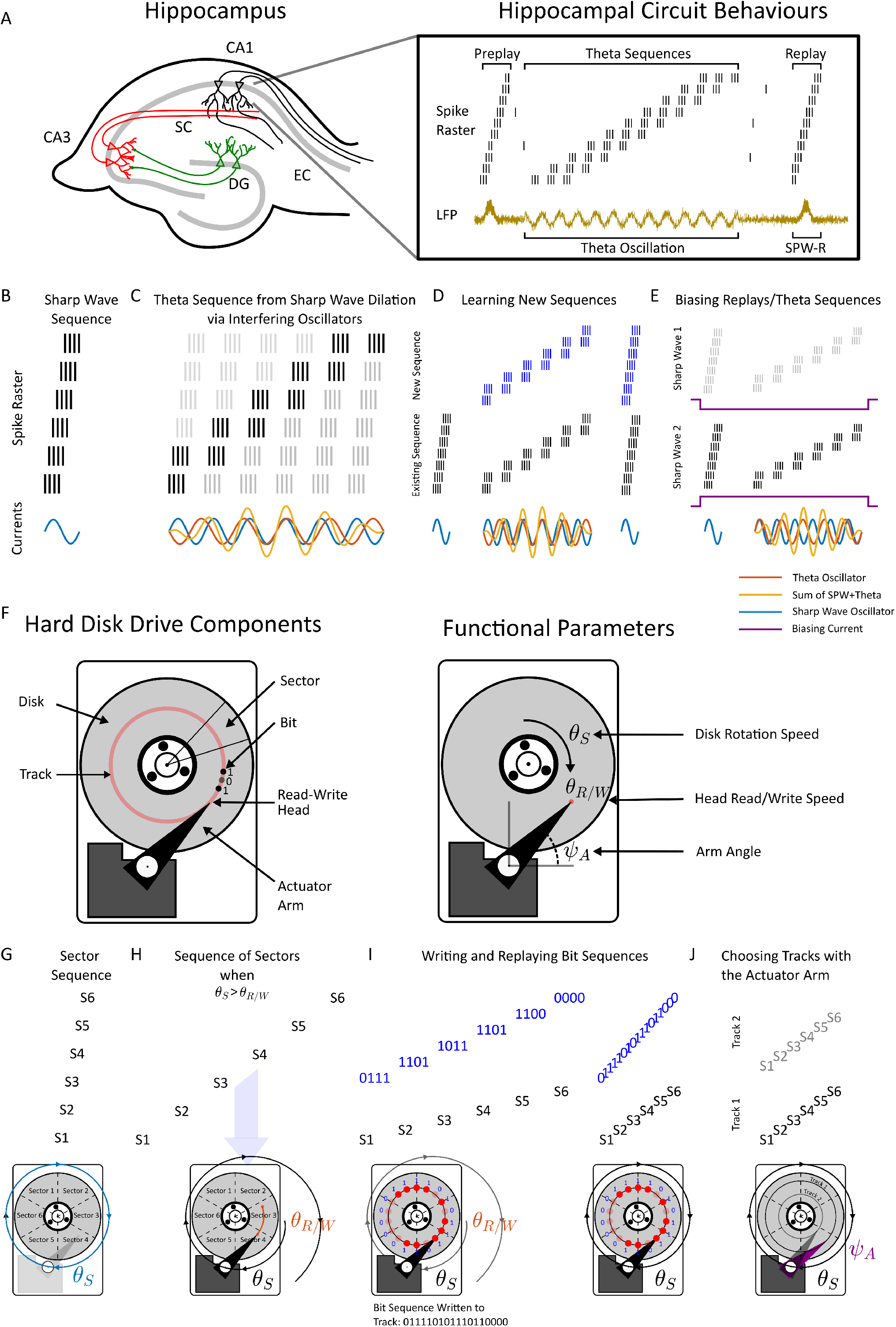
The links between hippocampal dynamics and disk drive operations. **(A)** The anatomical (CA3, CA1, Dentate Gyrus (DG), and Entorhinal Cortex (EC) sub-regions of the hippocanmpus (left) along with observed hippocampal behaviours (right). Sequences of spikes occur on short time scales as preplays/replays during hippocampal sharp-wave-ripples (SWRs), and on longer time scales as theta sequences. The components of a theta sequence during a theta oscillation are subsets of the entire preplay/replay. Preplays occur before the initial observation of a theta sequence, while replays occur after the initial observation of a theta sequence. The theta oscillation is an 8-12 Hz oscillation displayed in the local-field potential (LFP) while SWRs are 150-250 Hz high-frequency oscillations in CA1, which coincide with a large-deflection (the sharpwave) in CA3. **(B)** In a model of hippocampal replay, sharp-wave sequences are controlled by an intra-hippocampal oscillation. The oscillation is transiently activated for single cycles to trigger SWR replays. **(C)** By adding a secondary oscillation, SWR sequences become temporally dilated into theta sequences via an interference-based mechanism. **(D)** Dilated sharp-wave spike sequences can form a basis to learn new sequences, in a one-shot, instantly compressible format. **(E)** Different sharp-wave sequences can be elicited by a biasing current to pools of pyramidal neurons. **(F)** The hardware components (left) and functional parameters (right) of a computer disk drive, which operates by encoding data onto a rapidly spinning disk subdivided into tracks and sectors. The read/write head on the apex of the actuator armmoves to encode bits onto different tracks. The disk drive has three functional parameters: the Disk Rotation speed *θ*_*S*_, the read/write speed of the head *θ*_*R/W*_, and the actuator armangle *ψ*_*A*_.

We discovered that collectively, these operations could be explicitly mapped onto those performed by a computer’s Hard Disk Drive (HDD). To start constructing our brain-machine mapping of hippocampal dynamics, we first catalogued the individual components and operations of a disk drive and mapped these operations to the hippocampal equivalent (Figure 1F). The central operations of an HDD are performed by three components: an Actuator Arm (AA), a Rotating Disk (RD), and a Read/Write head (R/W) (Supplementary Material Section 1, Supplementary Figure S1, Supplementary Video 1, Figure 1E). The actuator arm points to a segment of the disk referred to as a track, which is further subdivided into sectors (Figure 1F). The read/write head, which is located on the apex of the actuator arm, writes new information in the form of bits, or reads previously stored bits on tracks and sectors. These three mechanical components are described by three dynamically evolving parameters that control the disk drive: the location of the actuator arm (*ψ*_*A*_, A for “Arm”), the disk rotation speed (*θ*_*S*_, s for “spinning”) and the read/write speed of the head (*θ*_*R/W*_, R/W for “read/write”, Figure 1F). As the disk spins in a single revolution (*θ*_*S*_), a sequence of sectors on a track appears directly beneath the R/W head on the actuator arm (Supplementary Video 1, Figure 1G). This sequence appears on the fast, intrinsic time-scale of the disk rotation speed (*θ*_*s*_). The sequence of sectors can be read from or written to in a single disk rotation (Supplementary Video 2, Supplementary Video 3). This mimics the compressed time-scale sequences which occur during hippocampal SWRs [1–8, 56, 73].

The sequence of sectors can also be accessed on a slower time-scale, by allowing the disk to perform slightly more than a full revolution for each read/write cycle of *θ*_*R/W*_ (*θ*_*R/W*_ *< θ*_*S*_ = *θ*_*S*_ − *ϵ*_*θ*_, Figure 1H, Supplementary Video 3). In this operating mode, the read/write head accesses each subsequent sector on a slower time scale which mimics the dilated or behavioural time-scale sequences that occur during hippocampal theta oscillations. Bit-sequences can be written to these sectors during the slower-access mode and then rapidly replayed after (Figure 1I, Supplementary Video 2). This is similar to the time-compressible, one-shot learning of spike-sequences observed in the hippocampus ([46, 48]). Finally, different tracks are accessible by the actuator arm when the actuator arm changes its location (Figure 1J). The location of the actuator arm, *ψ*_*A*_ behaves similarly to a line attractor. So long as no force is impinged on the actuator arm, the location *ψ*_*A*_ is constant and thus all actuator arm positions are stable. For an actual actuator arm, the range on *ψ*_*A*_ is restricted to some interval within [0, *π*]. Applying a force to the actuator arm moves *ψ*_*A*_ to a new track. This mimics how bias currents within the hippocampus, as a result of place, context, or other stimuli, can select different populations of cells during replays [73]. These biasing currents may be regulated or produced by attractor dynamics [74–78].

### 2.2 Constructing the Rotating Disk, Read/Write Head, and Actuator Arm Networks

Having qualitatively linked the dynamics of the hippocampus to those of a disk drive, we wondered if an explicit neural model could be constructed of the individual disk-drive components. Such a model, if constructed, would merge two modelling paradigms into a concrete device: interfering oscillators and attractor dynamics (see [79] for a complementary grid-cell model). To that end, we trained recurrent spiking neural networks to display the dynamics of a disk drive as a tripartite network with a rotating disk network, a read/write head network, and an actuator arm network [63, 80]. We started by modelling the three networks individually to reproduce disk-drive dynamics constrained to the hippocampal parameter ranges.

First, we considered the rotating disk network (Figure 2A-I). The RD network was trained with techniques in machine learning (FORCE training [80, 81]) to cycle in a sequence around a ring with frequency *θ*_*S*_ where *θ*_*S*_ was 9 Hz. This cyclical behaviour acts as the disk rotation of the NDD model with disk rotations mapping to SWRs (Figure 2A). A single isolated cycle around the ring corresponds to a single isolated SWR with a duration of approximately 100 ms (*θ*_*S*_^−1^). This rotational sequence is generated by asymmetrically coupling the interneurons in the rotating disk network on the ring while the interneurons are receiving a super-threshold excitatory current (Figure 2B). The interneurons that are currently firing in the sequence inhibit interneurons that have fired just before, thereby maintaining the sequential firing structure on the ring. This interneuron ring serves as the rotator for the entire disk. Subsets of pyramidal neurons serve as individual tracks in the rotating disk network. If the collection of neurons within a track receive enough excitation, they can fire unique sequences, commonly elicited during hippocampal SWRs (Figure 2C-D). The excitation comes from recurrent coupling between the pyramidal neurons with a strongly coupled subset of track initiators (Materials and Methods). The initiator neurons bind the subset of pyramidal neurons into a track and are activated stochastically (consistent with [82]) while the sequences within a track are regulated by the interneuron rotator. These initiators may be related to high-firing rate, low spatial specificity CA1 pyramidal neurons [83, 84].

**Figure 2:**
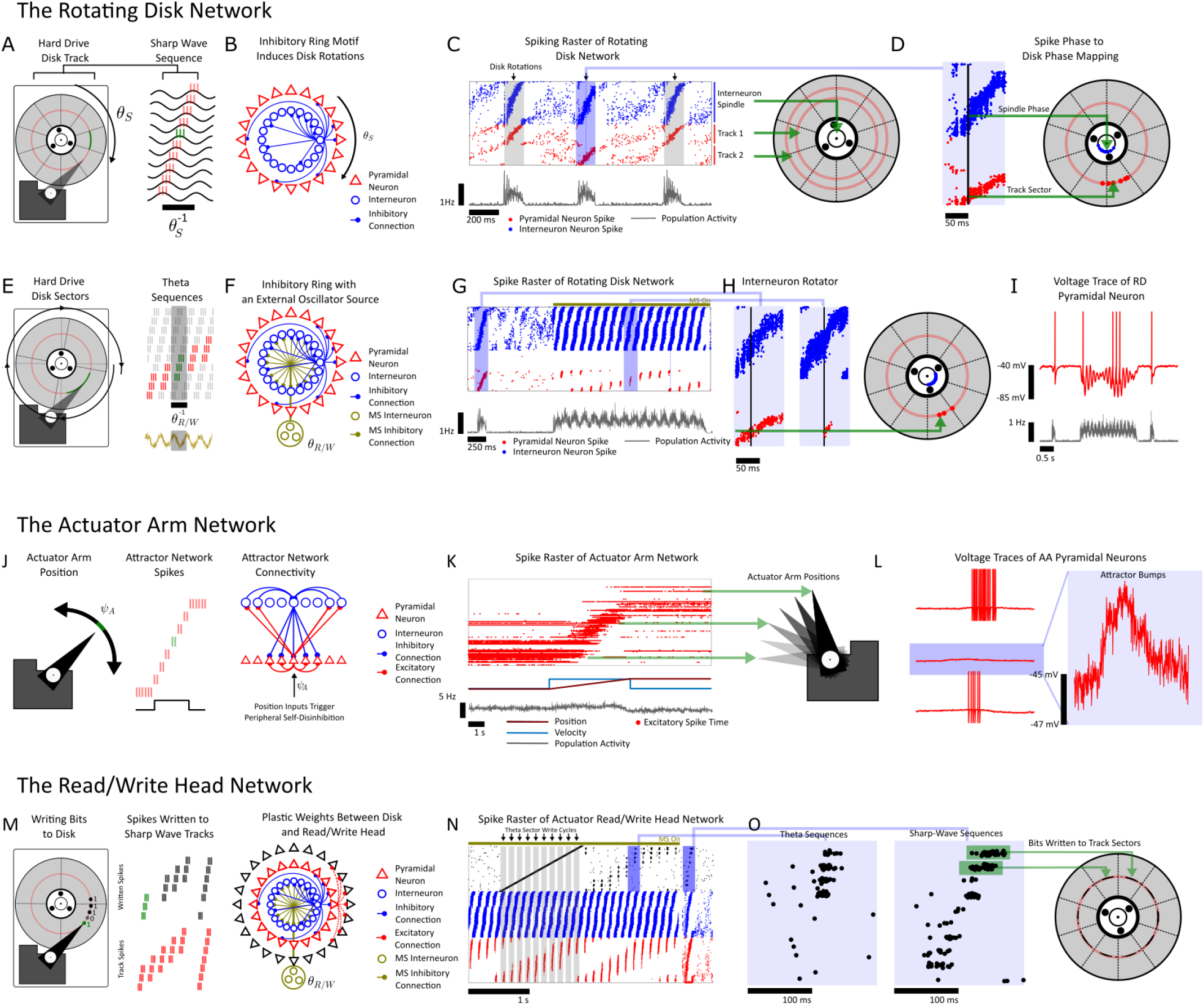
Mapping hippocampus circuit components onto hard disk drive components: the rotating disk, actuator armand read/write head networks. **(A)** The rotating disk (RD) network contains SWR sequences that instantiate tracks in the Neural Disk Drive (NDD) model of the hippocampus function. **(B)** An asymmetric ring of inhibition induces a rotating SWR sequence with rotational speed *θ*_*S*_. **(C)** Different subsets of RD excitatory neurons instantiate tracks in the NDD model. **(D)** The interneuron ring acts as the primary rotator of the NDD, while subsets of the RD SWR tracks act as the sectors. **(E)**The theta oscillations *θ*_*R/W*_ sequentially expose sectors of a track to write-to. This forms theta sequences, which in the NDD model, are sequentially exposed tracks of a SPR track. **(F)** An oscillatory inhibitory signal with a frequency of *θ*_*R/W*_ triggers sector exposure during a theta oscillation. **(G)** During a RD SWR, all sectors of a track are activated without *θ*_*R/W*_. During theta oscillations (with *θ*_*R/W*_), the sectors are sequentially activated. **(H)** Zoom of the spike sequences in the RD network overlaid explicitly onto a disk. **(I)** RD dynamics are present as theta oscillations in both the LFPs and in the membrane potential of individual pyramidal neurons. **(J)** The actuator arm(AA) features dynamics of an attractor network. A torque on the AA causes it to change angular position *ψ*_*A*_, which activates different subsets of AA neurons. This forms sequences of instantaneous rates, rather than individual spikes. When the inputs correspond to velocity and place, these ongoing firing fields can be interpreted as current place fields. **(K)** A simulation of the FORCE trained AA network with the position and velocity of the animal as inputs (below). The AA pyramidal neurons (red) are sorted according to their place preference. **(L)** The voltage traces for 3 randomly selected pyramidal neurons in the AA network. The voltage traces show “ramps of activity”, which allows the neurons to modulate their ongoing firing rates to collectively encode the position of the actuator arm. The position of the actuator arm can be decoded from a linear combination of these rates. **(M)** Once a track is selected by the AA, the read/write (R/W) head can write new sequences onto the track. This corresponds to updating the Schaffer collateral weights in the network to store new sequential information. **(N)** Spike raster plot for the RD and R/W head networks. The written sequences become discretized into assemblies of coactive neurons **(O)** The discretized assemblies projected onto the sectors of a disk.

Next, we found that pyramidal cells in the rotating disk network that constitute at track can be exposed sector(s)-at-a-time when a second oscillation with a slower frequency (*θ*_*R/W*_ = 8 Hz) is applied to the interneurons in the rotating disk network (Figure 2E). This frequency difference creates an interference that activates subsets of the full SWR-sequence of spikes (a track) in the rotating disk network but as subsets of a theta-sequence (Figure 2G-H). This is the slower operating mode where sectors can be accessed for reading/writing on a slower time-scale (Supplementary Video 3, Supplementary Video 4). We further observed that the interference between the *θ*_*S*_ and *θ*_*R/W*_ oscillations manifests as an interference pattern in the voltage of individual neurons (Figure 2) [85] and the interference produces internally generated theta sequences [25, 86, 87]. These internally generated theta sequences are used for reading or writing to tracks. Therefore, the rotating disk network accurately mimics both the operations of a spinning disk and the hippocampus as it consists of a series of pyramidal neurons arranged into tracks and sectors, which could be accessed on slow (behavioural) or fast (neural) time scales, and an interneuron ring which “spins” the entire disk as the rotator thereby forming spike sequences.

With the rotating disk network constructed, we then focused on modelling the actuator arm (Figure 2J-L). The actuator arm of a conventional disk drive operates much like a continuous line attractor. The actuator itself produces a physical force that moves the arm to a specified position, thereby selecting a track (Figure 2J). Once the force on the arm stops, the arm stays in its new position over the track and is stable. In that way, all possible positions of the actuator arm are stable while an applied force can rapidly switch the arm’s position, thereby forming a continuous line attractor.

Given the correspondence between an actuator arm and an attractor network, we trained a recurrent spiking network with FORCE training [80, 81] to mimic the dynamics of a line attractor [88] to serve as the actuator arm network (Supplementary Figure S2, Figure 2K). The actuator arm network receives two inputs: the desired position (*ψ*_*A*_) and velocity 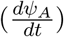 of the actuator arm. We trained the actuator arm network to integrate the velocity-like input to estimate the desired position 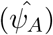. During training, the desired position, *ψ*_*A*_ was dropped during random periods to force the actuator arm network to produce a position estimate with integration alone (Supplementary Figure S2). After training, we found that the resulting actuator arm network produced isolated firing fields of pyramidal neurons, consistent with observed hippocampal place fields (Figure 2K-L, Supplementary Figure S3). However, here 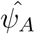 refers to the estimated position of the actuator arm on the disk, rather than the estimated position of an animal in physical space *per se*. Further, we observed that pyramidal neurons in the actuator arm network exhibited bumps of activity, reporting the dynamics of an attractor network (Figure 2L) [85]. Finally, we tested the line-attractor nature of the trained actuator arm network. When no velocity inputs and position inputs are applied, the actuator network retains its last known position as a stable state, similar to how a physical actuator arm stays in the last position on a real HDD if the actuator no longer provides a force on the arm (Supplementary Figure S4). Thus, the actuator arm network behaves as both a line-attractor network, and the actuator arm of a disk drive, while simultaneously producing hippocampal features such as place-field like firing in the individual cells.

Next, we wondered how the actuator arm network displayed its line attractor dynamics. To investigate this, we sorted all neurons (pyramidal and interneuron) according to their place (*ψ*_*A*_) preference (Supplementary Figure S3, Figure 2K). After sorting, we found that the actuator arm network maintained stable states through a self-disinhibitory motif. Pyramidal neurons with similar actuator arm position preferences excited both themselves and interneurons with similar *ψ*_*A*_ preferences. The interneurons then inhibited pyramidal neurons with dissimilar *ψ*_*A*_ preferences.

With the rotating disk and actuator arm networks created, we next investigated how information is written onto, and read from the hippocampal “tracks” in the rotating disk network network. For a physical disk drive, the read/write head on the actuator arm reads and writes bits of data (0’s or 1’s) on individual tracks as different magnetic field directions (e.g. ↑ or ↓) on a ferromagnetic material. We thus sought to identify how the read/write head network could “write” bits of information.

We thus mapped the functional capability of a R/W head in a disk drive to a plausible biological candidate: Hebbian plasticity (Figure 2D). We constructed a R/W spiking neural network that contains plastic excitatory synaptic weights from all pyramidal neurons in the RD network and uses these plastic weights to encode bits of information (Figure 2M). If enough excitatory weights from a track/sector connect onto a neuron in the R/W head, then that neuron spikes. The spiking of neurons in the read/write head (post-synaptic) paired with Hebbian plasticity to spikes in the rotating disk network (pre-synaptic) is how the read/write head network reads or writes bits of information.

Bits of information are sent to the R/W network from external sources for encoding, causing spiking in R/W pyramidal neurons. When the R/W neural spiking coincides with spiking from the active track in the RD network, Hebbian plasticity encodes bits of information in the AA→R/W pathway. The bits of information are written when the spinning disk continuously cycles (*θ*_*S*_), and the read/write oscillation (*θ*_*R/W*_) is active. As these oscillations sequentially expose the sectors of an individual track, bits are written discretely to isolated sectors (Figure 2N, Supplementary Video 3). We found that this discretization is visible as discrete assemblies when the information is subsequently replayed either when *θ*_*R/W*_ is active or in full disk rotations (Figure 2N-O, Supplementary Video 4) during SWRs, and in fact corresponds to the discrete sectors exposed during the theta-oscillation. This discretization of replays may be linked to the observation that replays can also “jump” from location to location discretely [89]. This result shows that the read/write head in the Neural Disk Drive Model can use Hebbian plasticity in a similar fashion to the read/write head on a physical HDD to encode bits of information onto a plastic medium. Finally, we remark that it may also be possible that different phases of the *θ*_*R/W*_ oscillation are used separately for the reading and writing of bits [90–93].

### 2.3 The Tri-Partite Neural Disk Drive Model

Next, we investigated if the rotating disk, Actuator Arm, and read/write head networks that we constructed separately could be assembled together and function synergistically to store information as the final Neural Disk Drive (Figure 3). We simulated these three constituent networks coupled with three sets of cross-network synaptic weights (Figure 3A). The first set of (AA→RD) connections link the actuator arm and rotating disk networks, selecting tracks and thus subsets of pyramidal neurons to access. The second set of (RD→R/W) connections associated the RD network to the R/W head network, and act as the storage media to store bits of information (Figure 3B). The third set of (AA→R/W) connections, from the AA network to the R/W network, triggers synfire-chain like spiking in the R/W network. This sequence of spikes represents the information to be acquired during the writing state/theta oscillation, and replayed during the reading stage/SWR. These three (AA, RD and R/W) networks acting together collectively operate as the hippocampal Neural Disk Drive (Figure 3C-F). The R/W head network served to record/replay information as spike sequences; that is, leveraging the computer analogy, sequences of bits are written/read onto a track (Figure 3C). The track is rotated by the ring of interneurons that mediates the disk dynamics (Figure 3D). The actuator arm network serves to perform path-integration and selects the specific track on the rotating disk network.

**Figure 3:**
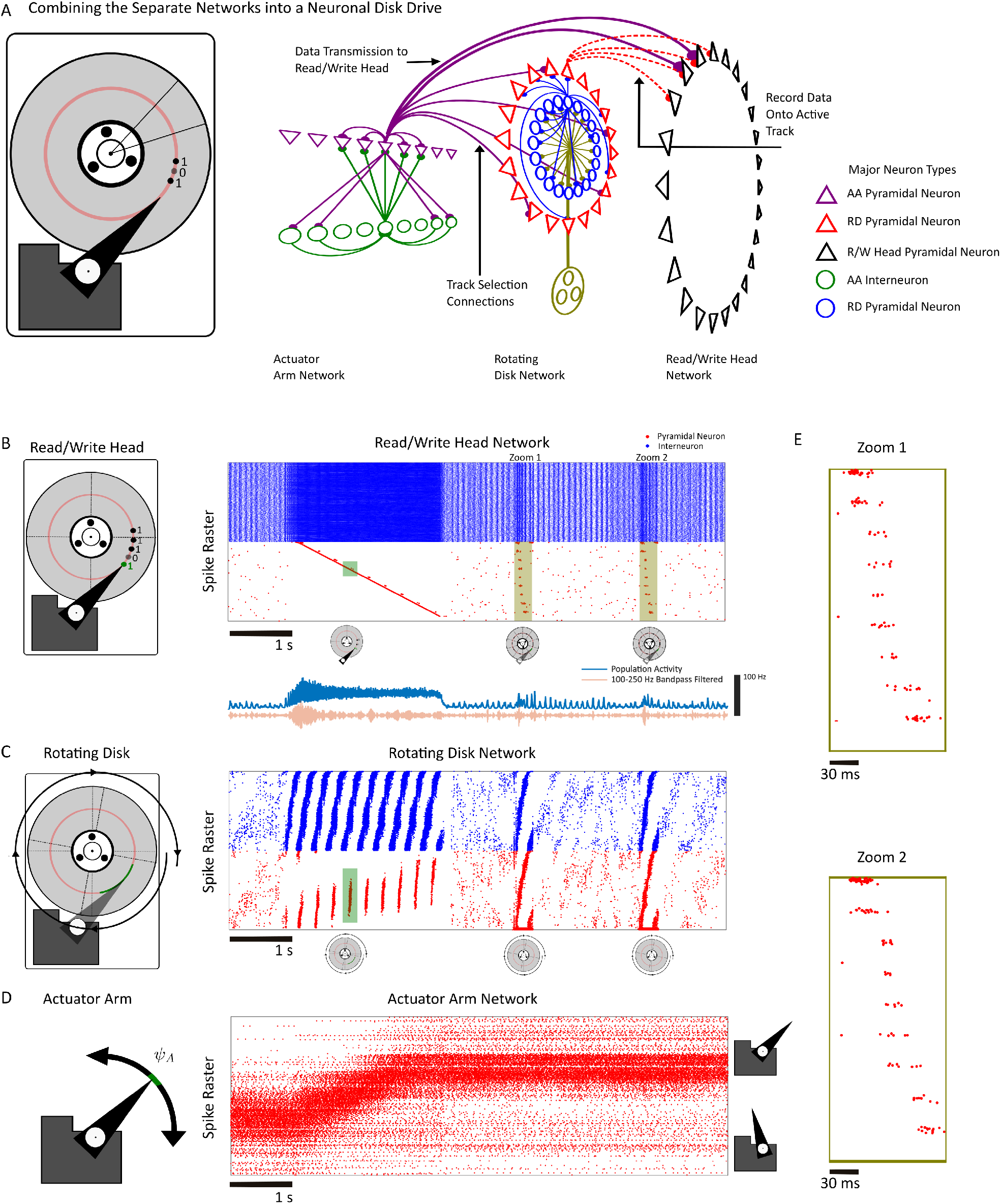
Assembling the tripartite Neural Disk Drive model. **(A)** The three constituent (RD, AA, and R/W) networks are connected together to construct a tripartite network. **(B)** The RD, AA and R/W networks correspond to discrete regions and pathways within the hippocampus circuit. **(C)** The R/W network is simulated with a writing component consisting of theta oscillations (*θ*_*R/W*_) and a reading phase during which SWRs occur (*θ*_*S*_). **(D)** The RD network during the writing phase and two disk rotations. The interneuron ring performs the disk rotations in both the writing and reading phase. **(E)** The AA network, with neurons sorted according to place preference. The AA transitions from one location to the next. **(F)** Two zooms of the replayed sequences that occurred during RD SWRs.

Together, these networks operate synergistically. For example, we found that the AA network can create (theta) sequences longer than those contained within a single track by switching between tracks (Supplementary Figure S5). The AA network can also bias which track becomes replayed during disk rotation (Supplementary Figure S5). Finally, the zoo of hippocampal data-accessing events are well explained by a disk-drive model of hippocampal operations. By disabling the rotating disk network interneurons, replays can be converted into sequence-less reactivations [94] (Supplementary Figure S6). Pre-plays, where the sequences during SWRs are observed before navigation [49–53,95], are pre-existing or old data written to tracks before they are accessed during Reading/Writing phases (Supplementary Figure S6). Fragmented replays [6], where replay trajectories jump in state-space correspond to mis-alignment between the initial phase of the disk and the start of a bit sequence (Supplementary Figure S6, Supplementary Video 5). Reverse replays based on dedicated sequence-reversion pools of interneurons were previously considered [63]. In this scenario, the disk spins backwards to reverse the order of bit/spike sequences (Supplementary Video 6). Replays of trajectories that were not previously experienced by an animal ([96, 97]) can also be constructed by reading out multiple tracks that were never sequentially accessed during the writing phase. These tracks can also be individually reversed within a multiple-track replay event [96, 97].

Thus, the Actuator Arm, rotating disk, and read/write head networks work synergistically to record information in a tripartite Neural Disk Drive, with core features of the hippocampal network being mapped to well-defined operations and components of a HDD.

### 2.4 Detecting the Interneuron Ring Sequences Predicted by the NDD Model

This Neural Disk Drive model proposes that the hippocampus uses both network oscillations and attractor dynamics to implement the operations performed by the rotating disk and the actuator arm of a computer disk drive. Thus, we tested this theoretical prediction using empirical data, probing evidence for one of the dynamical hallmarks of the NDD model: interneurons serving as the rotational backbone of neural firing patterns. To proceed, we first considered in vivo hippocampal ensemble recordings performed from rats trained to learn and remember three reward locations on a cheeseboard maze (Materials and Methods, Figure 4A). On each day of this memory task, animals develop an effective navigation path to reach the reward locations throughout the learning trials (Figure 4B). We used these animal trajectories to analyze hippocampal patterns formed by sequences of spikes (Figure 4C-D) with a Maximal Likelihood Estimation (MLE) based alignment algorithm (see Supplementary materials). The MLE algorithm aligns the repeating firing fields internally to each other across learning trials (rather than to place) by applying time-shifts across trials through each firing field (Figure 4D, Supplementary Figure S7). The time shifts are then used to optimize an objective function, the maximum-likelihood estimator for the spike density parameterized by said time shifts. We found that aligning a single neuron simultaneously aligns an entire community of neurons within the learning session (Figure 4E). This included both pyramidal neurons and interneurons (Figure 4F), unveiling a repeating interneuron sequence that appeared to be stable across learning trials (Figure 4G, Supplementary Figure S7). Interneurons in both mice and rats are known to have differing phase preferences with respect to the theta oscillation in the LFP. Indeed, we found a similar result (Figure 4H-K) with interneurons spanning the [0, 2*π*] range in both rats (Figure 4H-I) and mice (Figure 4J-K). However, phase preference firing of bursts in interneurons during theta oscillations, which has been previously reported ([98–100]) does not generate ring-sequences alone. Indeed, stochastic simulations of neurons with phase preferential firing of bursts in a population will not generate a ring sequence (Supplementary Figure S8). Our analysis shows that that hippocampal interneurons form the neural ring-sequences predicted by the NDD model across species and tasks.

**Figure 4:**
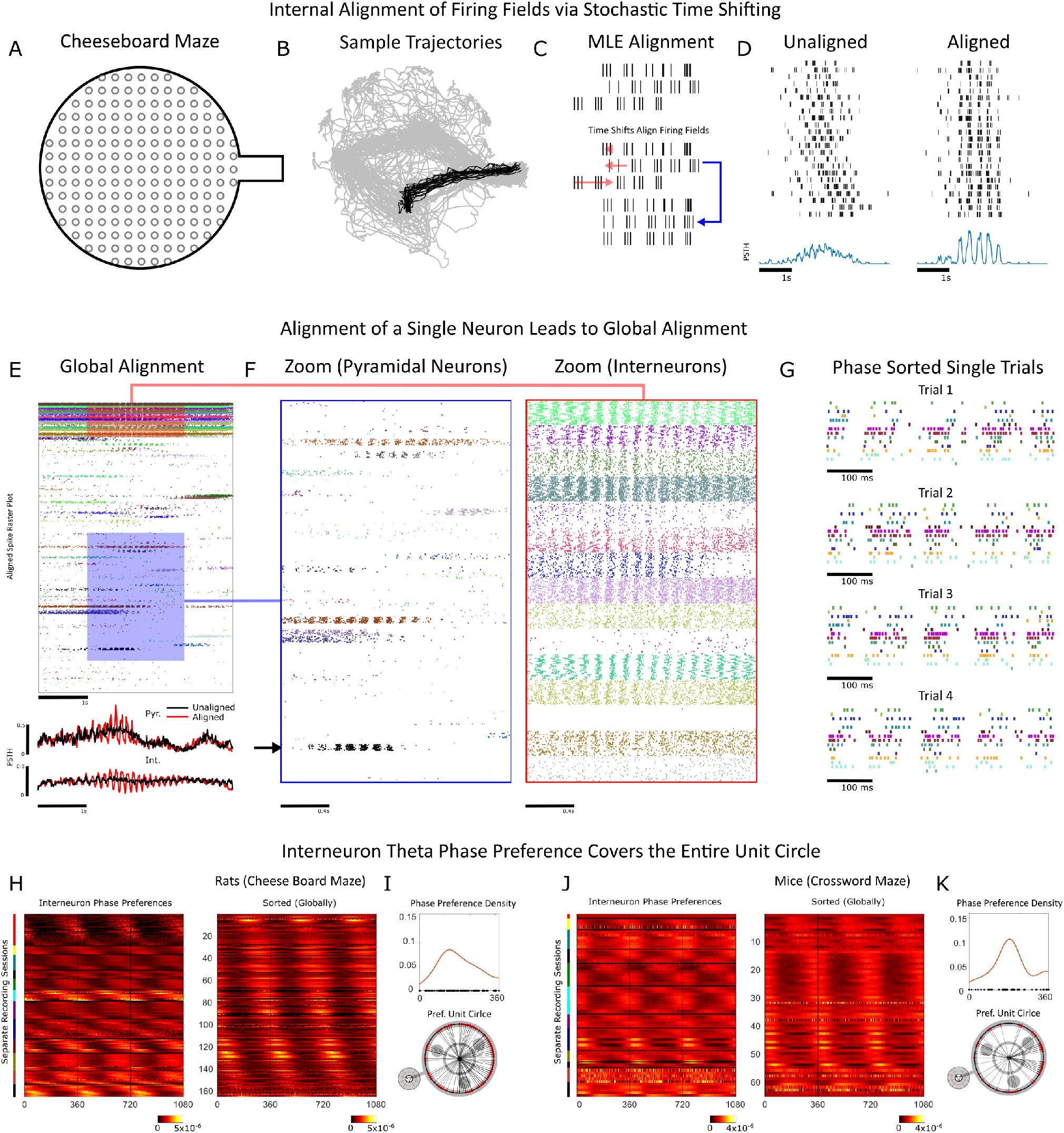
Probing the hippocampal in-silico NDD with in-vivo ensemble recordings. **(A)** Schematics of the cheeseboard and the crossword mazes where rodents (rats and mice, respectively) learn to navigate to reward locations while hippocampal neuron ensembles and LFPs are recorded. **(B)** Example repeated navigational trajectories (black) used for firing pattern analyses, superimposed on animal’s whole path (gray) in these mazes. **(C)** The trajectories are aligned with a maximum likelihood estimation (MLE) algorithm that maximizes the probability of recorded spiking with time-shifts to each temporal firing field. **(D)** The firing fields, temporally unaligned (left) and aligned (right). **(E)** Aligning a single recorded pyramidal neuron produces a global temporal alignment across the ensemble of multiple recorded neurons. **(F)** Zoom of the aligned recorded pyramidal neurons (left) and interneurons (right). The black arrow denotes the pyramidal neuron used to align the entire population. **(G)** A sequence of interneuron ring-like spiking. **(H)** Temporal firing fields for a population of interneurons. **(I)** The phase preference of firing for all interneurons is broadly distributed. **(J)** Similarly, the firing fields for a population of interneurons recorded in mice performing another memory task (the crossword maze). **(K)** The phase preference for all interneurons is once again, broadly distributed over [0, 2*π*].

## 3 Discussion

Analogies and metaphors with devices have been attempted towards obtaining useful descriptions of the brain since antiquity [14, 16–18]. Often, these analogies are a product of their times, manifesting brain function as a likeness or similarity to the dominant technological innovation of the era. Indeed, the computer analogy for brain function was proposed by Von Neumann shortly after both the first electronic computer, ENIAC (Electronic Numerical Integrator and Computer), was unveiled [101] and Hodgkin and Huxley successfully modeled the squid giant axon [102]. However, these brain analogies are often limited to observations of behaviour, and rarely directly linking the computations, components, and function of a device to the anatomy and physiology of any particular brain area [15]. This is where our contribution differs from the past by leveraging decades of in vivo experiments in the hippocampus.

We have constructed and simulated a model of hippocampal function as a Neural Disk Drive, by merging two prior modelling paradigms, attractors [74–78] and interference models [20] into one singular device. The network was constructed and simulated with 3 components: an actuator arm (Attractor), a rotating disk (Oscillator) and a read/write Head (for storing memories). This model was sufficient to reproduce some of the core behaviours of hippocampal neurons; path-integration and navigation, theta sequences and phase precession, compressed sequences and sharp-wave ripples, and the zoo of hippocampal replays. Finally, we verified one of the predictions of this model; a rotating ring-sequence of interneurons which constitute the rotator of the neural disk drive.

If the hippocampus does utilize oscillations and attractor dynamics similarly to how these dynamics are used in a disk drive, two natural lines of inquiry emerge. The first line of inquiry emerges from the saturation of tracks. A track on a physical disk drive can only hold so many bits. In fact, for a sufficiently large file, multiple tracks must be used to store the entire file (Supplementary Video 7). Thus, if the hippocampus stores information as a disk drive, does multi-track storage also occur? Evidence for the affirmative to this hypothesis is present in the literature in the form of so-called extended replays, where a trajectory is replayed as multiple, sequential sharp-wave-ripple complexes [28, 29]. These ripple complexes are separated by 150 ms intervals, with an inter-sharpwave-interval distribution displaying prominent peaks at multiples of 150 ms, hinting at a disk-rotation mechanism controlling SWR generation [63].

The second line of inquiry consists of the total storage space of the hippocampus, if it does indeed operate as a disk drive. In the NDD model, the storage medium used to write bits onto the RD network are the synaptic weights coupling the RD network onto the R/W head. Anatomically, these weights may be the Schaeffer Collateral connections linking CA3 to CA1, which constitutes a matrix containing at most *N*_*CA*3_*N*_*CA*1_ connections, where *N*_*CA*3_, *N*_*CA*1_ are the number of neurons in CA3, CA1. If we think of the weights as entirely binary, and consider reasonable values of *N*_*CA*3_ = *N*_*CA*1_ = *O*(10^5^) [103], we arrive at ≈ *O*(10^9^) bytes or *O*(1) GB of storage as an upper-bound for rats. Human estimates for *N*_*CA*3_ and *N*_*CA*1_ are larger (*O*(10^6^)), leaving *O*(10^2^) GB [104]. Thus, if the hippocampus does act as a disk drive, it is one of fairly limited storage, as 100 GB translates into roughly 100 hours of video and audio with 720p resolution. Additionally, these were upper bounds that were derived assuming all CA3 and CA1 neurons and that the neurons operate perfectly to transmit and store bits without redundancy. Given the low values of even these optimistic bounds, the neural disk drive model suggests that the hippocampus is limited in the amount of data it can store, possibly hinting that the hippocampus stores some type of data-compressed representation of complex memories [105–109]. Alternatively, it is also possible that the low amounts of storage are only used to store recent events, prior to consolidation [110–112].

## List of Supplementary Videos

To dynamically describe both the function of disk drives and the relation between hippocampal dynamics and disk drives, a series of animations were prepared (see below). All animations can be viewed and downloaded from https://www.nicolacomputationalneurosciencelab.com/publications.

**Supplementary Video 1: The Components and Operations of a Disk Drive**. A supplementary video that displays how the motions of the read/write head, rotating disk, and actuator arm stores and accesses information.

**Supplementary Video 2: Writing on the Slow Time Scale**. A supplementary video that shows how a disk drive can write with a frequency (*θ*_*R/W*_) that is slightly slower than the disk rotation speed (*θ*_*S*_). This frequency difference creates hippocampal phase precession in the Neural Disk Drive model.

**Supplementary Video 3: Replays via Single Disk Rotations**. A supplementary video that shows how a trajectory can be replayed in the Neural Disk Drive model with a single disk rotation.

**Supplementary Video 4: Theta Sequences via Multi-Sector Access**. A supplementary video that shows the emergence of theta-sequences from a tracks worth of data. Here, the read/write head accesses multiple sectors in a single disk rotation sequentially, thereby creating theta-sequences with phase precessing spikes.

**Supplementary Video 5: Fragmented Replay via Disk-Phase Misalignment**. A supplementary video showing how trajectories can “jump” via the misalignment of the initial phase or sector of the disk with the start of a trajectory.

**Supplementary Video 6: Reverse replay via Counter-Clockwise Disk Rotations**. A supplementary video showing how trajectories can be replayed backwards through time by spinning the disk in the opposite rotation (e.g. counter-clockwise) of the initial recording (e.g. clockwise).

**Supplementary Video 7: Extended Replay via Multi-Track Access**. A supplementary video showing how long trajectories can be decoded by using multiple tracks, with an actuator arm switching between successive tracks.

## Materials Availability

This study did not generate new unique reagents.

## Data availability

The datasets used in this will be made available via the MRC BNDU Data Sharing Platform (https://data.mrc.ox.ac upon reasonable request.

## Materials and Methods

### The Actuator Arm Network

The actuator arm network consists of 2000 excitatory (*AA*_*E*_), and 2000 inhibitory (*AA*_*I*_) leaky-integrate-and-fire neurons:

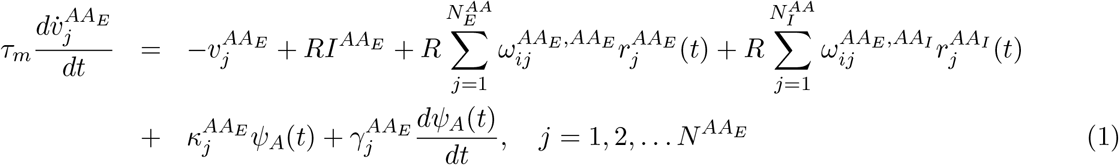

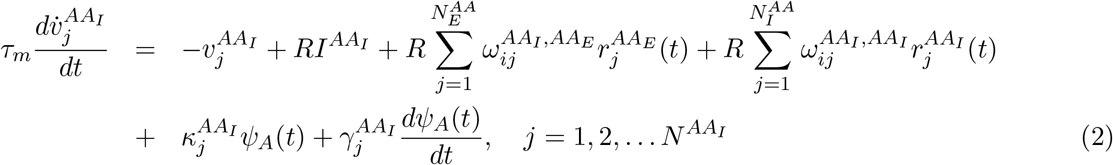

where 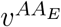 and 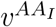 denotes the voltage for an actuator arm excitatory and actuator arm inhibitory neuron, respectively. The parameters for all neurons/synapses can be found in Table 1. Once the voltage for a neuron reaches a threshold, *v*_*thresh*_, the voltage is reset to *v*_*reset*_.

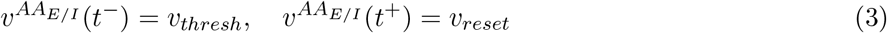

**Table 1:** The parameters used for the rotating disk (RD), actuator arm (AA) and read/write (R/W) Head networks, unless otherwise specified by the supplementary methods for each figure. The bias currents, *I*^*α*^, *α* = *RD, AA, R/W* vary to change the operational modes of the different sub-networks. Note that the nominal values of the bias currents may differ in specific figures/subfigures.

| Parameter Value | RD-E | RD-I | AA-E | AA-I | R/W-E | R/W-I |
| --- | --- | --- | --- | --- | --- | --- |
| $N$ | 2000 | 2000 | 2000 | 2000 | 4000 | 4000 |
| $t_{ref}$ | 2 ms | 2 ms | 2 ms | 2 ms | 2 ms | 2 ms |
| $t_m$ | 10 ms | 10 ms | 10 ms | 10 ms | 10 ms | 10 ms |
| $I^\alpha$ | | -25 pA/-40 pA | -40 pA | -40 pA | -41 pA/-42 pA | -40 pA |
| $v_{reset}$ | -65 mV | -65 mV | -65 mV | -65 mV | -65 mV | -65 mV |
| $v_{threshold}$ | -40 mV | -40 mV | -40 mV | -40 mV | -40 mV | -40 mV |
| $\tau_d$ | 20 ms | 20 ms | 20 ms | 20 ms | 20 ms | 20 ms |
| $\tau_r$ | 2 ms | 2 ms | 2 ms | 2 ms | 2 ms | 2 ms |

Every spike is followed by an absolute refractory period, *τ*_*ref*_ during which the neuronal dynamics are quenched at the reset value. The parameters 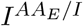, denote the bias currents to the neurons in the E/I sub-populations, respectively. The membrane time constant, *τ*_*m*_ controls the integration dynamics of each neuron. The parameter *R* = 1 · 10^9^ Ω serves as the resistance. The weight matrices 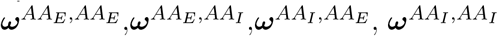 denote the coupling from *E* to *E, I* to *E, E* to *I* and *I* to *I* populations, respectively. These weights are trained with the FORCE algorithm, and described below. The inputs *ψ*_*A*_(*t*) and 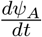 denote the desired position of the actuator arm, and the velocity of the desired position of the actuator arm. The inputs are multiplied by a set of input weights, 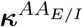, and 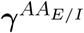 for the actuator arm position/velocity, respectively.

The variables 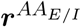 are the convolved spike times for the *E/I* actuator arm neurons:

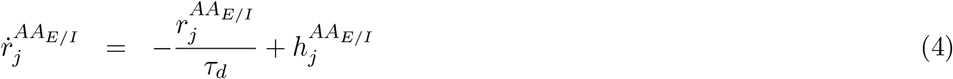

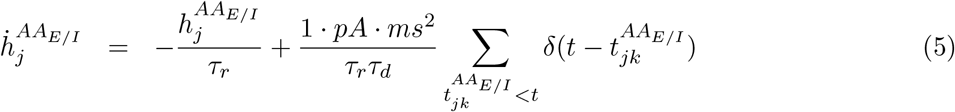

where 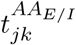 denotes the *k*th spike fired by the *j*th neuron in the actuator arm excitatory/inhibitory sub-population. The parameters *τ*_*d*_ and *τ*_*r*_ denote the decay and rise times, respectively of the sub-population of neurons. The AA network, and indeed, all networks considered were integrated with a simple Forward-Euler method and a step size of *dt* = 5 × 10^−5^*s*.

### FORCE Training the Weights of the Actuator Arm Network

The weight matrices 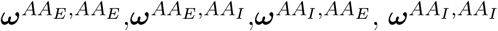 decompose as the sum of a static component, and a learned component:

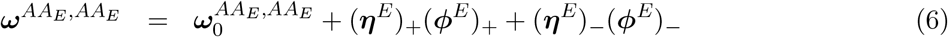

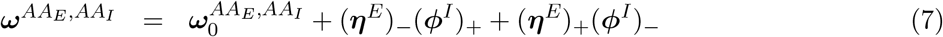

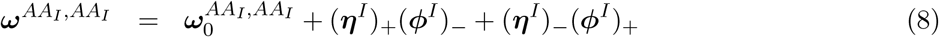

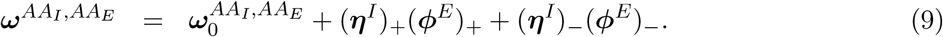

The functions

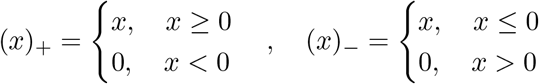

are applied to the components of the matrices ***η*** = [***η***^*E*^, ***η***^*I*^], ***ϕ*** = [***ϕ***^*E*^, ***ϕ***^*I*^] to enforce Dales Law. The matrices ***η*** are referred to as the encoders, and help determine the tuning properties of neurons with respect to the estimated actuator arm position 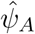. For each neuron in the AA network, the encoder for that neuron (a row of ***η***), is randomly generated and sparse. The encoder is an *N* × *k* matrix where *n*_*sup*_ is the dimension of the supervisor of the network, likewise for the decoder. The encoder is enforced to be sparse: each neuron has a single element in its encoder that is non zero, and randomly set to ±*W*, where *W* = 10pA. The input weights, 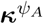 and ***κ***^*γ*^ were randomly generated with from a uniform [−1, 1] distribution, for all neurons in the AA network.

The decoders, ***ϕ*** are learned with FORCE training, which we describe below.

#### The Supervisor and Inputs to the Actuator Arm Network

The input to the actuator arm network is a randomly generated signal, *ψ*_*A*_(*t*), and its derivative, 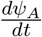. The signal is generated with a bounded, double-filtered noisy process. The first filter corresponds to the acceleration *a*(*t*) while the second filter corresponds to the velocity *ψ*_*A*_(*t*) :

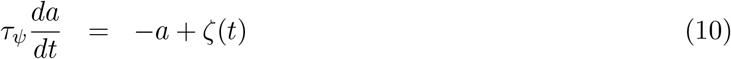

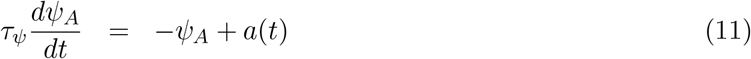

where *ψ*_*A*_(*t*) = ± 1 causes a reset, as if colliding with a boundary, to the velocity and acceleration of the actuator arm position 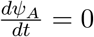, *a*(*t*) = 0. Further, the velocity of the actuator arm is also limited such that if 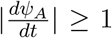, the velocity is fixed to ±1, to prevent arbitrarily fast motion of the actuator arm. The variable *ζ*(*t*) is a white noise process with mean 0 and standard deviation of 10^−3^. The actuator arm position is unit-less, while the velocity and acceleration are *s*^−1^ and *s*^−2^.

The position and velocity inputs 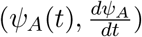 are provided to the actuator arm network during training. However, the position component is dropped stochastically (set to 0) for intervals that are randomly generated. These intervals are a minimum of 1 second long, and a maximum of 51 seconds long, with the interval itself drawn from the uniform distribution *U* ([1, 51]). Once the position dropping interval ends, the position is turned back on instantly, for a random period of time. This random interval with position is also uniformly generated from the distribution *U* ([1, 51]). In the intervals where the position is dropped, the network must rely on velocity and the last known position of the system to “integrate” and estimate the desired actuator arm position.

The supervisor, ***s***(*t*) to the actuator arm network is a *n*_*sup*_ = 200 dimensional vector that is a non-linear transform of the position:

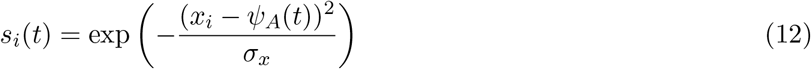

Each component of the supervisor acts as an activity bump when *ψ*_*A*_(*t*) passes near *x*_*i*_, where *x*_*i*_ is the center of the bump. The centers are uniformly distributed on the interval [−1, 1]. The variable *σ*_*x*_ controls the width of the bump, with *σ*_*x*_ = 0.3 used during training. The velocity component is not contained in the supervisor in any way.

#### Recursive Least Squares

The decoders, ***ϕ*** are determined dynamically to minimize the squared error between the approximant and intended dynamics, ***e***(*t*) = ***ŝ***(*t*) − ***s***(*t*). The Recursive Least Squares (RLS) technique updates the decoders to solve this problem in real-time:

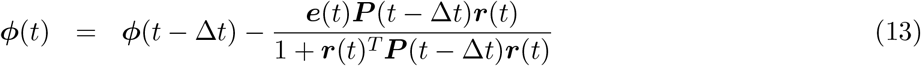

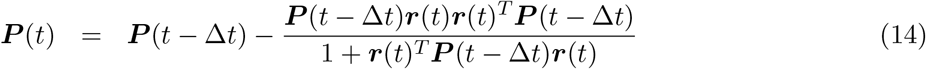

and 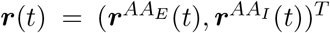. The Recursive Least Squares Algorithm and FORCE training is described in greater detail in [80, 81]. The actuator arm network is initialized with ***ϕ***(0) = **0, *P*** (0) = ***I***_*N*_ *λ*, where ***I***_*N*_ is an *N* -dimensional identity matrix, and *λ* controls the learning rate of RLS. The value *λ* = 0.5*dt* was used, where *dt* = 5 × 10^−5^*s* was the simulation integration step size. To implement Dale’s law as in equations (6)-(9), we decompose ***ϕ*** into ***ϕ***^*E*^ and ***ϕ***^*I*^ with ***ϕ***^*E*^ = (***ϕ***)_+_ and ***ϕ***^*I*^ = (***ϕ***)_−_. The training parameters for the RD network. For the rotating disk network, a value of *λ* = 0.05*dt* was used.

### The Rotating Disk Network

The rotating disk network is a modification of the so-called “SHOT-CA3” network from [63]. As in the actuator arm network, the rotating disk network consists of coupled leaky integrate-and-fire neurons:

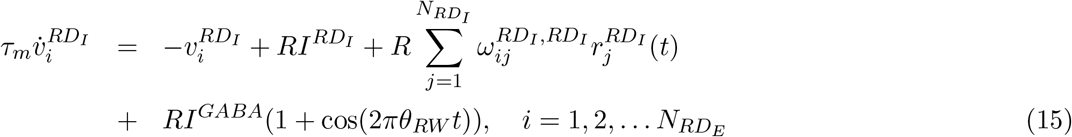

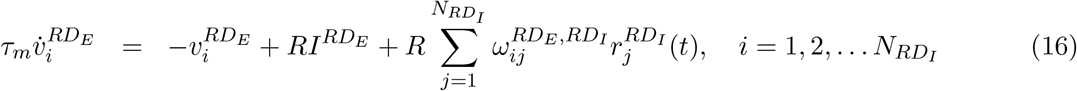

where *RD*_*E*_, *RD*_*I*_ denote the excitatory and inhibitory populations of the rotating disk network. The neurons receive a constant background current *I*^*α*^ for *α* = *RD*_*E*_, *RD*_*I*_. The *RD*_*I*_ neurons receive an oscillatory input where *θ*_*RW*_ is the input frequency, and *κ >* 1 determines the tonic level of inhibitory drive. The INP-MS has amplitude *I*^*GABA*^ = −10 pA for *i* = 1, 2, … *N*_*I*_. The network consists of 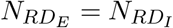 spiking neurons.

The weight matrices 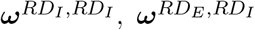 and supervisor used to generate them with FORCE training are described in further detail below, and in the specific methods for individual figures. They decompose similarly to the actuator arm network weights in equations (6)-(9). The supervisor used to train the RD network is a bank of oscillators:

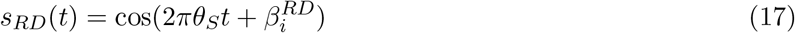

where *ϕ*_*i*_ is a uniformly distributed [0, 2*π*] phase preference for each oscillator. The phase preferences 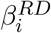 are randomly drawn from a uniform distribution on the interval [0, 2*π*]. All weight matrices that we consider are dimensionless with the units of current (pA) carried by the synaptically filtered spike trains ***r***(*t*) (see Equation (5)).

### The Read/Write Head Network

The read/write Head network is a population of leaky integrate-and-fire 2000 inhibitory (*RW*_*I*_) and 2000 excitatory (*RW*_*E*_) leaky-integrate-and-fire neurons:

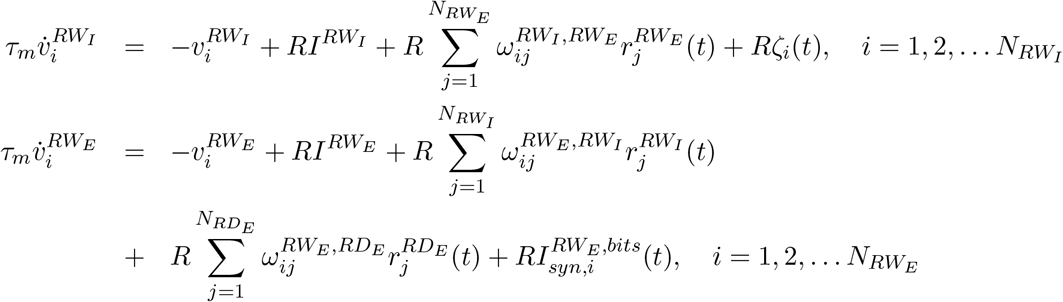

where *ζ*_*i*_(*t*) is an independent white noise term with mean 0 and standard deviation *σ* = 0.2 pA. This noise term prevents the pathological synchronization of interneurons. The weight matrices 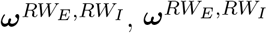 are untrained, and described below. The bits sent to the *RW*_*E*_ neurons are expressed as time dependent currents 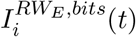. Finally, the weight matrix 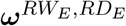. The matrix 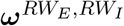 is a random matrix with each element drawn from a uniform distribution 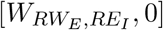 where 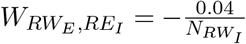. The matrix 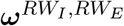 is also randomly generated, on the interval 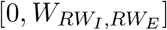 where 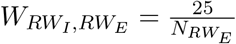.

The weights from the rotating disk excitatory neurons to the read/write head excitatory neurons are learned with a Hebbian-plasticity based learning rule ([63]):

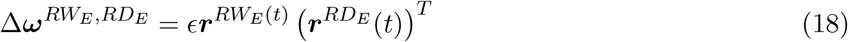

For efficiency in the numerical simulations, the update rule (18) is applied every 15 *dt* time steps, rather than every time step. The parameter *ϵ* acts as a learning rate for the synaptic weight adjustments and controls how rapidly the weights are adjusted.

## Maximum Likelihood Alignment

### Identifying Repeating Trajectories

To maximally align the spike times, we first selected a trajectory component from the rats navigating the cheeseboard maze from [38], which was restricted to 4 seconds in duration and contained large, linear movements along the maze within those 4 seconds. The initial selected trajectory ***p***_0_ = (*x*_0_(*t*), *y*_0_(*t*)), for *t* ∈ [*t*_0_, *t*_0_ + 4] was then used as a motion template, where other ***p***_*j*_ = *x*_*j*_(*t*), *y*_*j*_(*t*), *t* ∈ [*t*_*j*_, *t*_*j*_ + 4] were found by proximity to the initial template via the *L*_2_ norm:

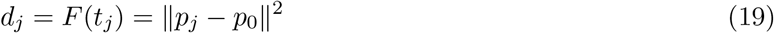

The trajectory components were found by treating *t*_*j*_ as a continuous variable, *τ* and floating *τ* over the entire interval, *τ*:

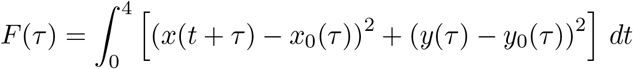

Then, a peak detector algorithm (*findpeaks*, MATLAB 2020a), was used to detect local minima in *F* (*τ*) (maxima in -*F* (*τ*)). Only the top 70% of these peaks were used, as *F* (*τ*) may contain local minima that are dissimilar from the initial trajectory *x*_0_. The set of minima of *F* (*τ*), correspond to the discrete times 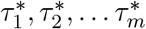.

### Performing the Maximum Likelihood Alignment

With the trajectory alignment times 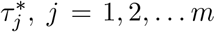 determined, a single neuron was selected for target alignment. The spikes for the *m* trials of that neuron were selected, and a kernel density estimator was constructed, *ρ*_*ner*_(*t*),where *t* ∈ [0, 4]. The bandwidth of the Gaussian kernel was taken to be 5 milliseconds. Then, for the *m* trials, a random *m* × 1 vector of *m* time shifts was generation. Each scalar component of this vector would would shift all the spikes within one of the *j* = 1, 2, … *m* trials by a constant amount, 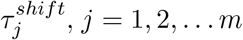. The goal of these random time-shifts is to determine the time-shift vector ***τ*** which would minimize the following quantity, commonly referred to as the cross entropy:

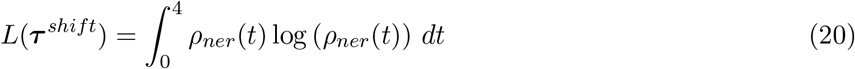

Minimizing the cross entropy is mathematically equivalent to maximizing the log-likelihood-function with the vector of shift times ***τ***^*shift*^ serving as the parameters. This procedure is commonly referred to as Maximum-Likelihood Estimation (MLE) of parameters. We remark that alternative methods can also be used to align the spike times [113].

To minimize the cross entropy, we employed an iterative stochastic gradient descent based algorithm. At every time point, the best shift vector, 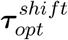 so far is perturbed:

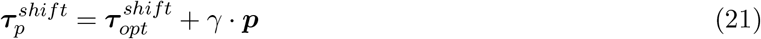

where *γ* is a variable learning rate, and ***p***_*n*_ is an *m* × 1 normally distributed random vector, from a standard normal distribution. The optimal time shift is then updated as:

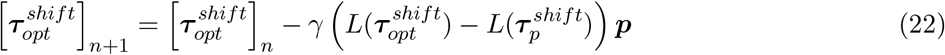

which serves to estimate the gradient of *L* as a function of ***τ***^*shift*^ and descend the gradient in one-step. This stochastic gradient algorithm is run for *n* = 10^4^ iterations for all animals, with an initial *γ* = 10^−1^. Every 10^3^ time steps, *γ* is halved. This slows down the learning rate of this stochastic algorithm for longer times, and yields more precise solutions to 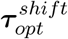.

With the MLE alignment parameters 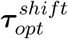 determined for a single neuron on [*t*_*j*_, *t*_*j*_ + 4], for the *j* = 1, 2, … *m* trials, the spike times are then shifted by the *j* the component 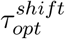. The 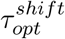 determined for aligning a single neuron is used for all neurons in the spike-raster plot with spikes son [*t*_*j*_, *t*_*j*_ + 4].

## Specific Methods for Figures/Supplementary Figures

Figure 2

In Figure 1E-F, the rotating disk network is simulated for a total time of 10 seconds with the *θ*_*RW*_ oscillator input off. For Figure 1I-K, the *θ*_*RW*_ oscillatory input is turned on for 2 seconds in the interval [3, 5]s. All neurons in the rotating disk network additionally receive a white noise current with mean 0 and standard deviation 0.5 pA, to mimic the stochastic firing/SWR initiation observed during slow-wave sleep. The background currents to all neurons, except to the initiators were −40 pA, while the initiator neurons were set to −40.1 pA. Larger or smaller currents to the initiators controls the SWR average rate, while larger/smaller currents to the *RD*_*I*_ neurons controls both the SWR average rate, and the shape of the inter-SWR-interval distributions [63]. For all simulations in Figure 1, the *RD*_*E*_ neurons were split into two tracks. The tracks were assigned by first constructing a random permutation of the rows of 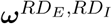, splitting the permutation into two sets, and then sorting the permutation with respect to the *θ*_*S*_ phase preference. This is mathematically equivalent to splitting the population of *RD*_*E*_ neurons into two sets, and sorting them on the unit circle. Each track had 50 initiator neurons, and 950 neurons representing the sectors (phases) of the disk. The initiators in each track connect to all other intra-track initiators with a strong recurrent weight 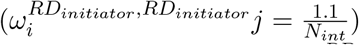, and connect to all other excitatory track neurons, and rotating disk neurons with the weight 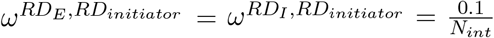. To implement inter-track competition, the excitatory neurons within a track project to rotating disk interneurons with randomly drawn weights from the interval 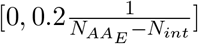. In Figure 1I, we biased one of the tracks towards being preferentially activated by increasing the bias currents to the track neurons, similarly to the impact the actuator arm network would have. For Figure 1I, In the reading mode, all initiators in Track 1 had a bias of −40.3 pA, while those in Track 2 had a bias of −40.5 pA. All remaining track neurons had a bias of 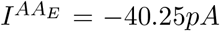 and 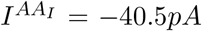. All excitatory neurons in all tracks and initiators also have a spike-frequency adaptation variable, *u*_*i*_(*t*), which increases by an amount of *d* = 18 pA for every spike fired by that neuron and decays with a time constant of 50 ms. The adaptation current is negatively weighted, and serves to slow down or even eliminate repetitive spiking. This adaptation variable can stop a disk-rotation/sharp-wave-ripple. In the writing-mode/theta-oscillation mode, the bias currents for all non-initiator excitatory neurons is increased to −6*pA*. For Figure 1K-L, the total simulation time is 25 seconds, with a non-zero, constant velocity (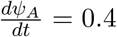 given in the interval [10, 15] seconds. The *AA*_*E*_ in Figure 1K neurons were sorted according to their phase preference, as determined in Supplementary Figure 3. For clarity, only a subset of *AA*_*E*_ neurons (10%) are plotted. In Figure 1N, the R/W head network is simulated for 15 seconds, with 5.1 seconds in the write mode. The information written to the network is a synfire-chain of spikes elicited in the read/write head neurons by an external current. The external current is an additional 20 pA applied to each neuron for 40 ms, in sequence. In the read/SWR-mode, the bias currents to the read/write head neurons are 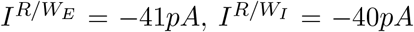. In the write/theta-oscillation mode, the bias currents are unchanged for the R/W neurons, while 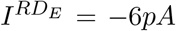 and 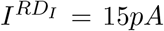. For both the read and write modes, the bias current for the initiators was −40.3 pA.

Figure 3

The actuator arm, rotating disk, and read/write head networks were combined into a single network with a 10 second long simulation time. The actuator arm network was identical to all previously considered (Figure 2, supplementary figures) implementations. The signal actuator arm position was generated as the integral of a constant signal, 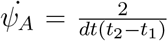, over the interval [*t*_1_, *t*_2_] with 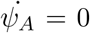 outside this interval. The desired actuator arm position is initialized with *ψ*_*A*_(*t*) = −1 for *t < t*_1_. This initialization results in a linearly ramping *ψ*_*A*_ from −1 to 1 on [*t*_1_, *t*_2_], and *ψ*_*A*_ = 1 for *t*_2_ *>* 1. The values *t*_1_ = 1, and *t*_2_ = 2.2 were used. The rotating disk was as in Figure 1, only with a single track and a single population of initiators. The intra-initiator weight was also stronger, 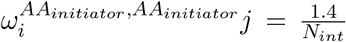, which was necessary to generate replays in the read/write head. The adaptation step size was also increased to *d* = 22*pA* in all rotating disk pyramidal neurons, to terminate sharp-wave bursts. The actuator arm projects random, sparse excitatory connections to all non-initiator neurons in the rotating disk network. The weights are drawn from a uniform distribution with mean 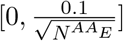 with probability *p* = 0.1 and 0 otherwise. The actuator arm also projects to a set of 500 leaky-integrate-and-fire neurons which act to generate the synfire chain of spikes, which serves as the bits to be written in this example. This network has identical parameters as all other neurons, only with a chain of connectivity between these neurons to trigger the synfire chain of spiking. Each neuron in the chain is connected to the next with a synaptic weight of 0.2. These neurons receive a strong hyperpolarizing bias current of −60.09*pA*. A 20 pA current pushes these cells in the superthreshold firing regime where the current is slightly over-threshold (−40.09 pA). All other bias currents are identical to prior implementations in Figure 2 for the isolated networks in reading and writing modes. The learning rate, *ϵ*, was 1.2 × 10^−5^.

Figure 4

The spikes in figure 4H-K were mapped onto LFP phases, with the LFP taken as be the channel with the most spikes detected. The raw signal was further band-pass filtered with a butterworth filter with a [4, 12] Hz window. The phase from the LFP was computed first via a Hilbert transform, and then the application of the arctangent function (*atan2* in MATLAB 2020a). Only epochs of theta oscillations, as defined in [38] were used in our final analysis. All spikes were then transformed into phases via a linear interpolant (*interp1* in MATLAB 2020a). The spikes were subsequently duplicated from [0, 2*π*] into the intervals [2*π*, 4*π*] and [4*π*, 6*π*], with a simple histogram with binds of width 6*π/*100 used to compute the spike-base histograms on [0, 6*π*]. The neurons were subsequently ordered in sequence according to increasing mean-phase. An identical protocol was applied to the rat data also.

Supplementary Figure 2

The total simulation time is 400 seconds, with the first second of simulation time used to initialize the chaotic spiking neural network. Recursive Least Squares (RLS) is turned on for the next 350 seconds and subsequently turned off. The last 49 seconds of simulation time are used for testing the network performance. The network was initialized in the balanced inhibitory regime: 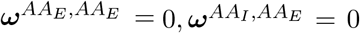, that is, all initial excitatory weights were set to 0, and all excitatory weights were learned. This initial state lets us constrain the firing rates of pyramidal cells to arbitrarily low rates. The bias currents were 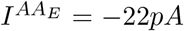, and 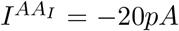. The initial inhibitory weights were randomly generated with exactly 90% sparsity. Each of the inhibitory neurons made 200 connections, with the connection strength set to 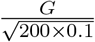, where *G* = −0.1. We found that this was sufficient to initialize the network into a chaotic spiking (prior to learning), and therefore serve as an adequate reservoir for RLS. RLS was only applied every 20 time steps (Δ*t* = 20) for efficiency, as the weight updates in equations (13)-(14) are *O*(*N* ^2^) in time complexity.

Supplementary Figure 3

The trained AA network was simulated for 200 seconds, with a randomly generated actuator arm position *ψ*_*A*_(*t*) and 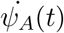, generated similarly to the initial training supervisor. The actuator arm signal, *ψ*_*A*_ remained on for all 200 seconds to establish the neural preference of firing to the *ψ*_*A*_ coordinate. The locations corresponding to each spike times were determined with a linear interpolant (*interp1*, MATLAB 2020a). For each neuron, a histogram was constructed with 41 bins of size 0.05, distributed uniformly over the [−1, 1], which is the operating range of *ψ*_*A*_(*t*). For each *AA*_*E*_ and *AA*_*I*_ neuron, the maximum of the spike-*ψ*_*A*_ histogram was determined. The maxima were sorted in ascending order (Figure SF 3A), and the AA neurons were re-ordered in an identical fashion to expose the weight structures (Figure SF 3B-C). The bias currents were 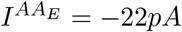, and 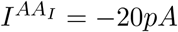.

Supplementary Figure 4

The trained AA network was simulated for 50 seconds, with a randomly generated actuator arm position *ψ*_*A*_(*t*) and 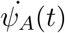, generated similarly to the initial training supervisor. The bias currents were 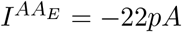, and 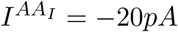. The position input *ψ*_*A*_ was turned off (set to 0) in Figure SF 4D, while the velocity input was turned off in Figure SF4E after 25 seconds. In all three simulations, the same random initial seed is used to generate *ψ*_*A*_ and 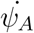.

Supplementary Figure 5

For both simulations, the rotating disk network was subdivided into two tracks with 50 initiators each. In the reading/sharp-wave mode, the bias currents were 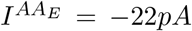, and 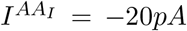, and 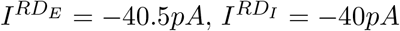. The initiators had a bias current of −40.7 pA. In the writing/theta-oscillation mode, the BIAS currents were the same for the actuator arm networks, while 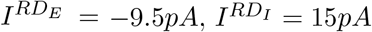. The initiators in the writing mode were kept off with a hyperpolarizing current (−60*pA*). In the reading mode, the initiators received connections from the AA neurons. The weights were given by the scaled, FORCE-trained decoder, 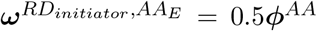 for the actuator arm position variables *ŝ*_*i*_(*t*). The first tracks 50 initiators received 10 duplicates of the first 5 components of *ŝ*_*i*_(*t*), while the second tracks initiators received 10 duplicates of the last 5 components of *ŝ*_*i*_(*t*). In the writing/theta oscillation mode, the track neurons in the rotating disk network received scaled decoders as weights, 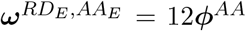. In this case, the 200 components of ***ϕ*** were duplicated 5 times each, scaled up, and provided to the rotating disk network as inputs from the actuator arm excitatory neurons.

Supplementary Figure 8

The neuron consisted of 20 Poisson neurons which generate spikes stochastically as part of an inhomogeneous Poisson process. First, in each theta cycle, the individual neurons were modeled as having a phase preference in generating bursts with the preference given by:

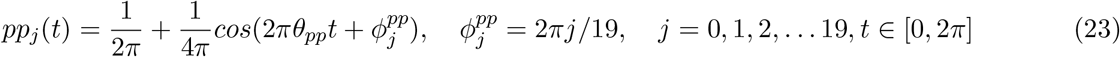

where *θ*_*pp*_ = 8*Hz*. In each cycle, a cell is allowed to fire a single burst with a phase-centre drawn from the probability distribution *pp*_*j*_(*t*). The phase centre for the *j*th neuron on the *i*th cycle, *χ*_*ij*_ is then used to generate a burst of spikes with probability:

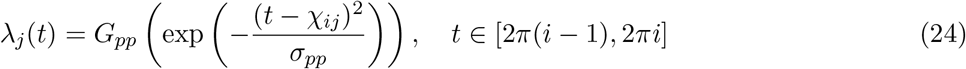

where *σ*_*pp*_ = 0.1 controls the width of a burst, and *H*(*x*) is the Heaviside function, and *G*_*pp*_ = 0.4 sets the spikes emitted per burst. The value of *G*_*pp*_ was selected so that approximately 10 spikes per burst were emitted. As measured empirically, the cells fired on average 13.11 spikes per cycle, which was estimated after 2000 cycles of firing.

## Supplementary Materials

Supplementary Section S1: Disk Drive Dynamics

A Hard Disk Drive (HDD) is a computer component dedicated to storing information. The simplest type of HDD consists of two primary sub-components: A rotating disk and an actuator arm (Figure S1A, [47]). The actuator arm moves across the disk to different regions as the disk is spinning at a constant rotational velocity. The disk is spun by a central spindle, which we will refer to as the “rotator” in the main text to avoid confusion with *sleep spindles*. The head of the actuator arm then writes information onto the disk by magnetizing a layer of ferromagnetic material. The magnetic field imprinted on the disk can be oriented in different directions, such as up or down, thereby allowing the encoding of binary information or bits as these magnetic field directions. The head of the actuator arm can then read data from the disk after the writing procedure is complete. For a fixed position of the actuator arm, the region on the disk that the head can access (through disk rotation) is called a track or cylinder (Figure S1A).

The physical components of the HDD give rise to 3 parameters that describe its function: the disk rotation speed *θ*_*S*_, the head writing speed *θ*_*W*_, and the angle of the actuator arm *ψ*_*A*_ (Figure S1B). The disk rotation speed is the number of revolutions per second that the disk makes, while the heard writing speed is the number of bits per second that the disk writes at. Finally, the actuator arm angle dictates which track the head writes information to. Thus, disks operations can be summarized by three parameters: two oscillation frequencies and a position variable for the actuator arm. These variables map nicely onto the dominant models of hippocampal function: dual oscillators and attractor networks.

For a standard HDD, we expect that the write speed is significantly faster than the disk rotation speed (*θ*_*R/W*_ ≫ *θ*_*S*_, Figure S1C). In this regime, each write cycle of *θ*_*R/W*_ takes place when the disk has advanced to the next sector on the track. Thus, the exact sequence of bits is maintained and encoded onto the disk on the corresponding sequence of sectors. Then, after writing has concluded, the sequence of bits in a single track can be read in a single revolution of the disk (Figure S1D). Thus, hard drives preserve the sequence of bits, and maximize the number of bits written to a track by ensuring that the write speed is significantly faster than the disk rotation speed. We will refer to this as the Hard Drive “nominal” parameter regime, as this is how a typical HDD stores information.

However, the sequence of bits can still be encoded with slower writing speeds. In particular when the disk writing speed (*θ*_*R/W*_) is slightly slower than the disk spinning speed (*θ*_*S*_), *θ*_*R/W*_ = *θ*_*S*_ − *ϵ*, sequences of bits can still be written to a track (Figure S1E). In this regime, the disk makes slightly more than a full revolution in between write cycles. This ensures that the head writes to the next sector of the track, even under the constraint that the writing speed and disk rotation speeds are similar. Once again, a single revolution of the disk can subsequently read all the information in a track. We refer to this operating range, when *θ*_*R/W*_ = *θ*_*S*_ − *ϵ* for a small frequency difference *ϵ* as the Hard Drive “precession” parameter regime. Typical HDDs do not operate in this regime.

When the writing speed is slightly faster than the rotation speed (*θ*_*R/W*_ ≈ *θ*_*R/W*_ +*ϵ*), the disk makes slightly less than a full revolution in between write cycles. This implies that rather than writing to the next sector of the track, the head writes the next bit to the previous sector of the track (Figure S1F). The effect of this operating regime is to encode information in the reverse order that it was observed. A single revolution in a read cycle can then read the entire bit sequence in reverse. We refer to this as the Hard Drive “recession” parameter regime. As in the HDD precession regime, typical HDDs do not operate in this regime.

Finally, when the head has saturated a track with a sufficiently long sequence of bits, the only recourse to store more information is to change *ψ*_*A*_ and encode more information onto new tracks (Figure S1G). Thus, multiple read cycles are now necessary to read out sequences that can not fit on a single track (Figure S1H). In particular, the disk must complete multiple full rotations, with the actuator arm switching between tracks in each rotation.

**Supplementary Figure 1:**
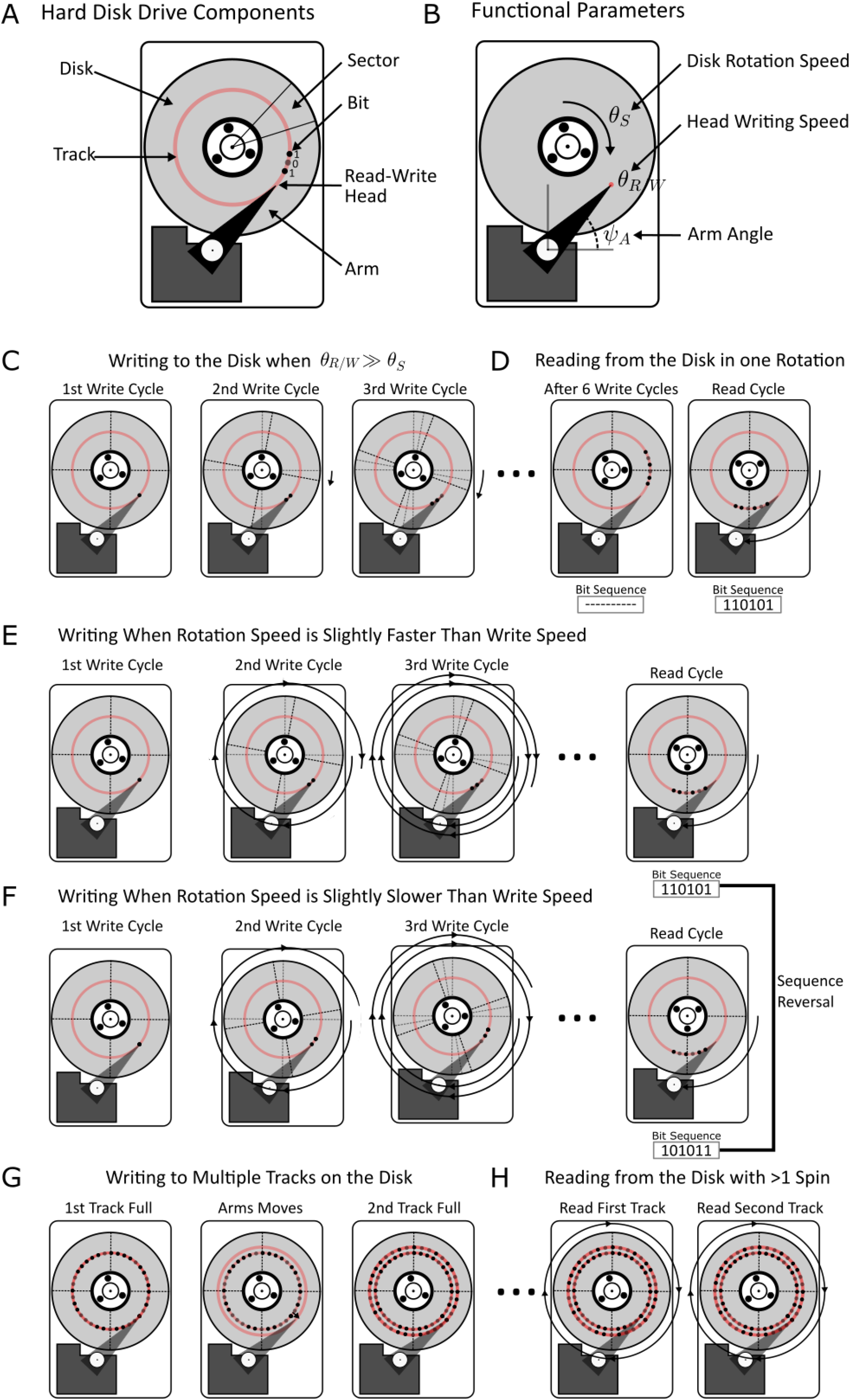
The Operations of a Hard Disk Drive **(A)** A Hard Disk Drive (HDD) schematic with its basic components. The rotating disk is used to store information onto tracks, which correspond to a circular segment of the disk. A sector on the disk is a wedge shaped region formed by any two radii. The actuator arm points to discrete tracks, which are the regions of the disk a stationary actuator arm has access by virtue of disk rotation alone. Bits are encoded onto the rotating disk by a read/write head on the apex of the actuator arm. The bits are encoded by inducing magnetic fields of different directions onto a ferromagnetic material on the disk surface. **(B)** The functional parameters of an HDD that describe its state: the disk rotation speed (*θ*_*S*_), the angular position of the actuator arm, *ψ*_*A*_, and the head writing speed, *θ*_*R/W*_. **(C)** When the write speed *θ*_*R/W*_ is substantially faster than the disk rotation speed, bits are written continuously as the disk spins. **(D)** Written bits on a single track can be read in a single revolution of the disk by the read/write head. **(E)** Bit sequences can also be written when *θ*_*R/W*_ = *θ*_*S*_ − *ϵ*, where the writing to a track corresponds to slightly greater than a full revolution of the disk. The written information can be read in a single disk rotation, as in **(D). (F)** When the disk rotation speed is slightly slower than the writing speed, the disk has advanced slightly less than a full revolution when a bit is written. This results in the bits being encoded and readout in the reverse order with which they were encoded. **(G)** When a track is saturated with bits, the actuator arm must move to the next track on the disk to write additional information. **(H)** Long sequences of bits must be written to multiple tracks. In order to access this information, multiple disk rotations are required with the actuator arm adjusting between tracks. This results in multiple read cycles/disk rotations required to readout information.

**Supplementary Figure 2:**
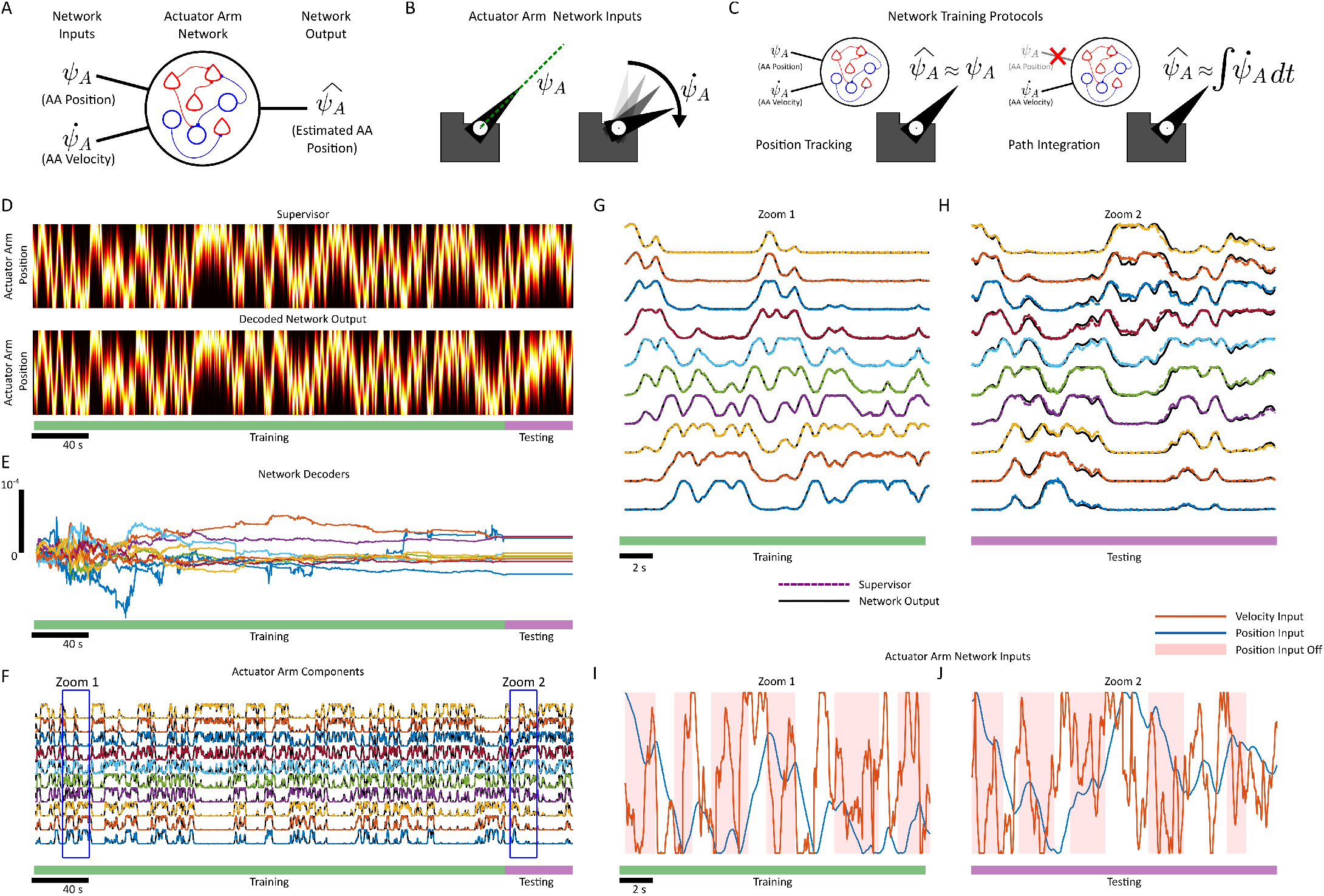
FORCE Training the actuator armNetwork **(A)** A network of 2000 leaky integrate-and-fire pyramidal neurons and 2000 leaky-integrate-and-fire interneurons is collectively trained to estimate the position of the actuator arm, 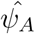, on a disk drive. **(B)** The network receives two inputs, a position variable *ψ*_*A*_, and a velocity variable 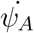. **(C)** The position variable and velocity variables can set the position of the actuator arm in the actuator armnetwork. When both inputs are on, the network outputs an actuator arm position 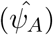 as determined by its position input *ψ*_*A*_. When the position input is off, the network is trained to integrate the velocity input 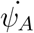. **(D)** The supervisor used to train the actuator arm(AA) network consists of 100 components that encode actuator arm position, forming a high-dimensional supervisor [80]. The components are sorted according to the supervisor-encoding preferences with respect to the actuator arm position *ψ*_*A*_. As a result, the supervisor forms a bump of activity, indicating the actuator arm position 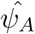. The training signal is randomly generated, with the actuator arm position being randomly dropped during training. FORCE training was as applied for 230 seconds (green), after the first second of simulation time to remove any transients. FORCE training was turned off in the last 19 seconds (purple) of the simulation. The supervisor (top) and decoded network output (bottom) are shown as heat maps. **(E)** A subset of the decoders, ***ϕ***_*i*_, *i* = 1, 2, … *N* during (green) and after (purple) FORCE training. (**F**) A subset of the 100 supervisor components from (A)-(B). The network output is plotted as coloured dashed lines, while the supervisor components are in solid black. **(G)** A 10 second zoom of subset of the supervisor and network components while the FORCE training is on. **(H)** A 10 second zoom of the subset of the supervisor and network components while FORCE training is off. **(G)** The inputs (AA velocity in blue, AA position in orange) to the network during training. The red shading denotes the position input was turned off during this time period. **(H)** The inputs (AA velocity in blue, AA position in orange) to the network after training. The red shading denotes the position input was turned off during this time period.

**Supplementary Figure 3:**
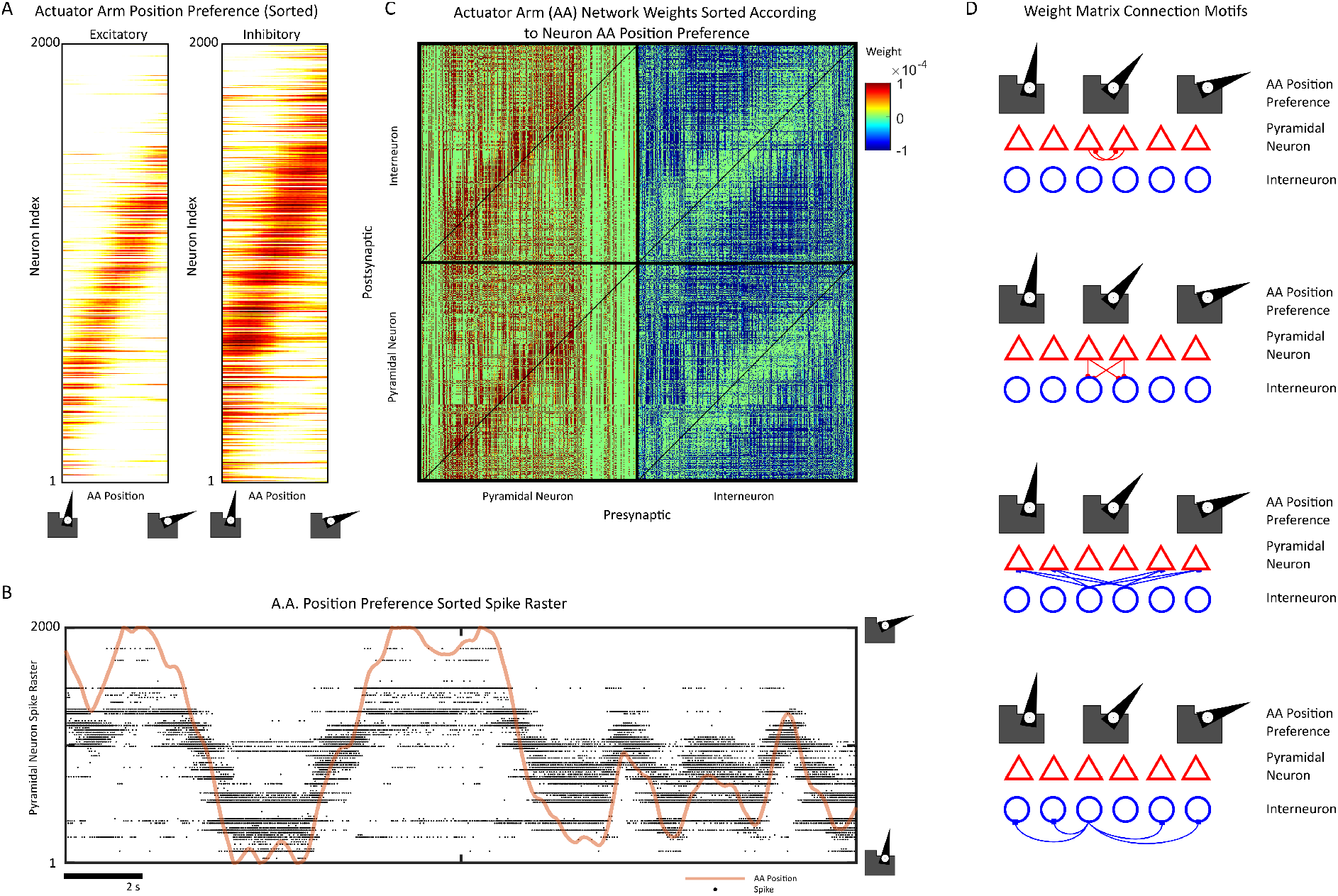
Weight Structure of the Force Trained actuator armNetwork **(A)** The 2000 pyramidal neurons (left) and 2000 interneurons (right) are plotted in order of their firing preference with respect to the actuator arm position *ψ*_*A*_. The preference is estimated by binning the spikes into a histogram on the actuator arm range *ψ*_*A*_ ∈ [−1, 1]. Both interneurons and pyramidal neurons display firing preferences with respect to *ψ*_*A*_, although the pyramidal neuron preferences are narrower. **(B)** A spike raster plot of the pyramidal neurons sorted according to the firing preference in *ψ*_*A*_. The position is scaled and overlaid for comparison. **(C)** The *ψ*_*A*_-preference sorted weight matrices for the pyramidal and inhibitory neurons. Excitatory connections, which are exclusively made by pyramidal neurons are plotted by red, while inhibitory connections, are plotted in blue. All *EE, EI, II*, and *IE* weight matrices show clear banding along the main diagonal. Note that the green vertical band is caused by a sub-population of pyramidal neurons that did not fire strongly during navigation, and as a result, do not display strong place-preferences and thus continue the banding structure. These neurons were also sorted to the top 400 pyramidal neurons due to the MATLAB *sort* function. **(D)** The four connection motifs created by FORCE training and shown in the banding structure. Pyramidal neurons tend to excite pyramidal neurons with similar *ψ*_*A*_ preference. Pyramidal neurons excite interneurons with similar *ψ*_*A*_ preference. Interneurons inhibit pyramidal neurons with different *ψ*_*A*_ preference. Interneurons inhibit interneurons with different *ψ*_*A*_ preference.

**Supplementary Figure 4:**
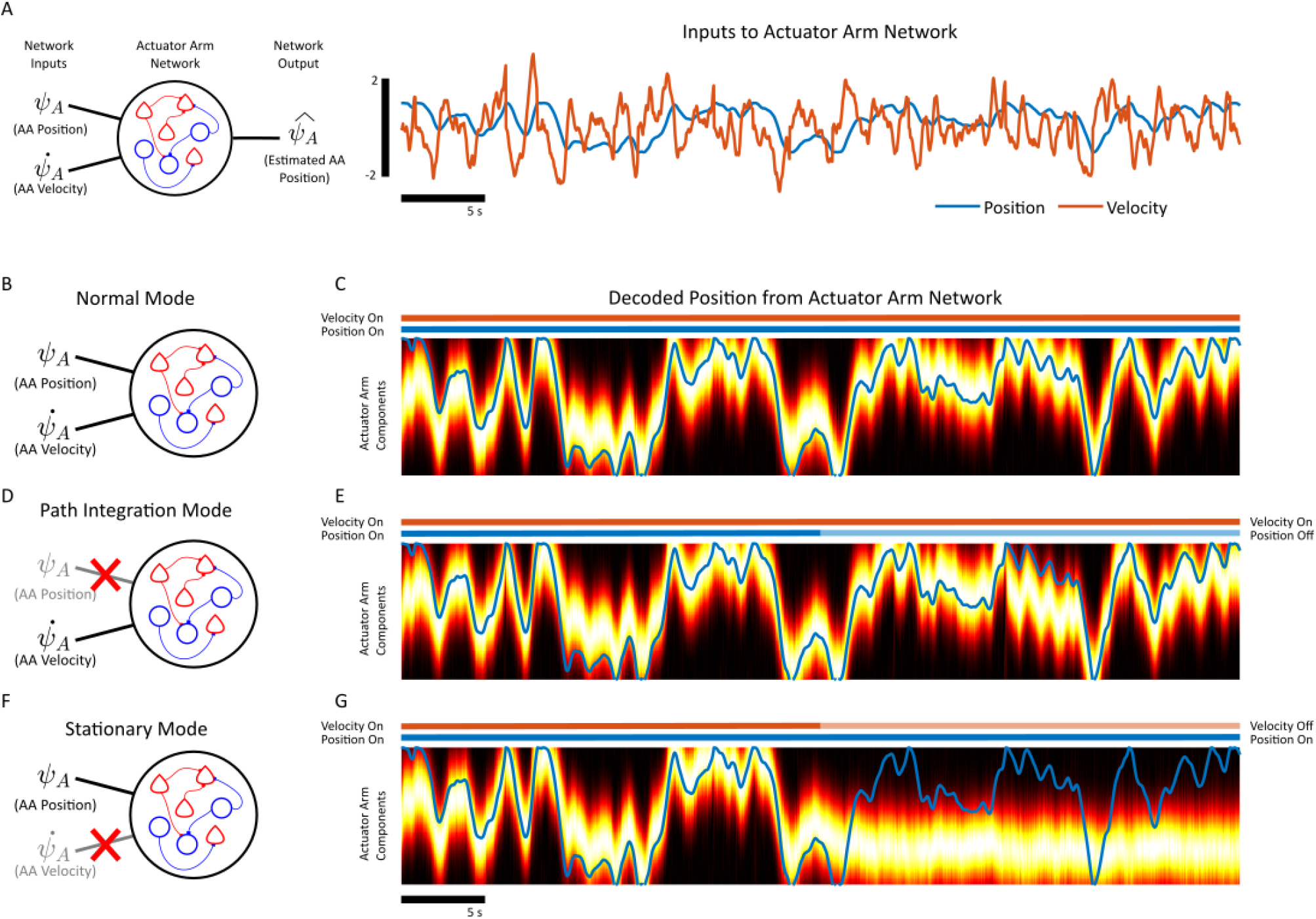
Operating Modes of the actuator armNetwork **(A)** The actuator arm network receives two signals, a velocity signal 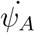 (orange) and a position signal *ψ*_*A*_ (blue). The position signal denotes the desired position of the actuator arm. The velocity signal is randomly generated, while the position signal is the integral of the velocity signal. **(B)** Under “normal operating mode”, the position signal and velocity signal are both applied. **(C)** The position of the actuator arm 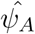 closely tracks the desired position *ψ*_*A*_ for a 50 second simulation of the AA network **(D)** In the path integration mode, the position signal *ψ*_*A*_ is not present. **(E)** The position signal is dropped in the last 25 seconds of simulation. The actuator arm network still tracks the desired position of the actuator arm by integrating the velocity signal 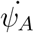. **(F)** In the stationary mode, the velocity signal 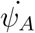 is dropped while the position signal is still applied. **(G)** The velocity signal is dropped in the last 25 seconds of the simulation. The network interprets this drop as a velocity of 0, and as such, retains an approximation of the last known position of the actuator arm.

**Supplementary Figure 5:**
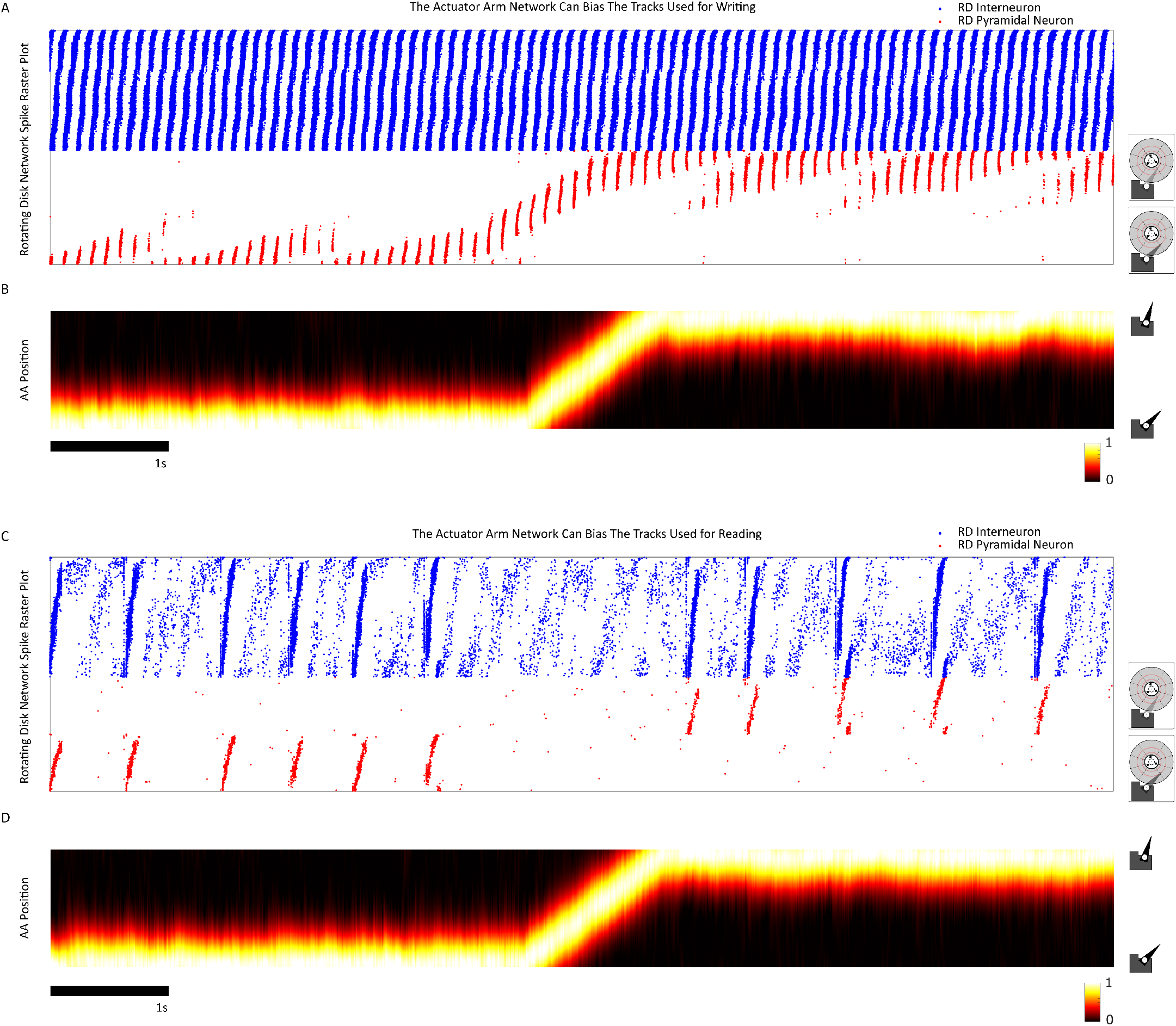
The actuator arm network can select tracks on the rotating disk Network **(A)** The spike raster plot for the excitatory (red) and inhibitory (blue) spikes for the rotating disk network while both oscillations *θ*_*S*_ and *θ*_*R/W*_ are operating. The rotating disk network is separated into two tracks with 950 neurons each. **(B)** The decoded position of the actuator arm. **(C)** The spike raster plot for the rotating disk network when only the *θ*_*S*_ oscillation is transiently induced by noise. **(D)** The position of the actuator arm.

**Supplementary Figure 6:**
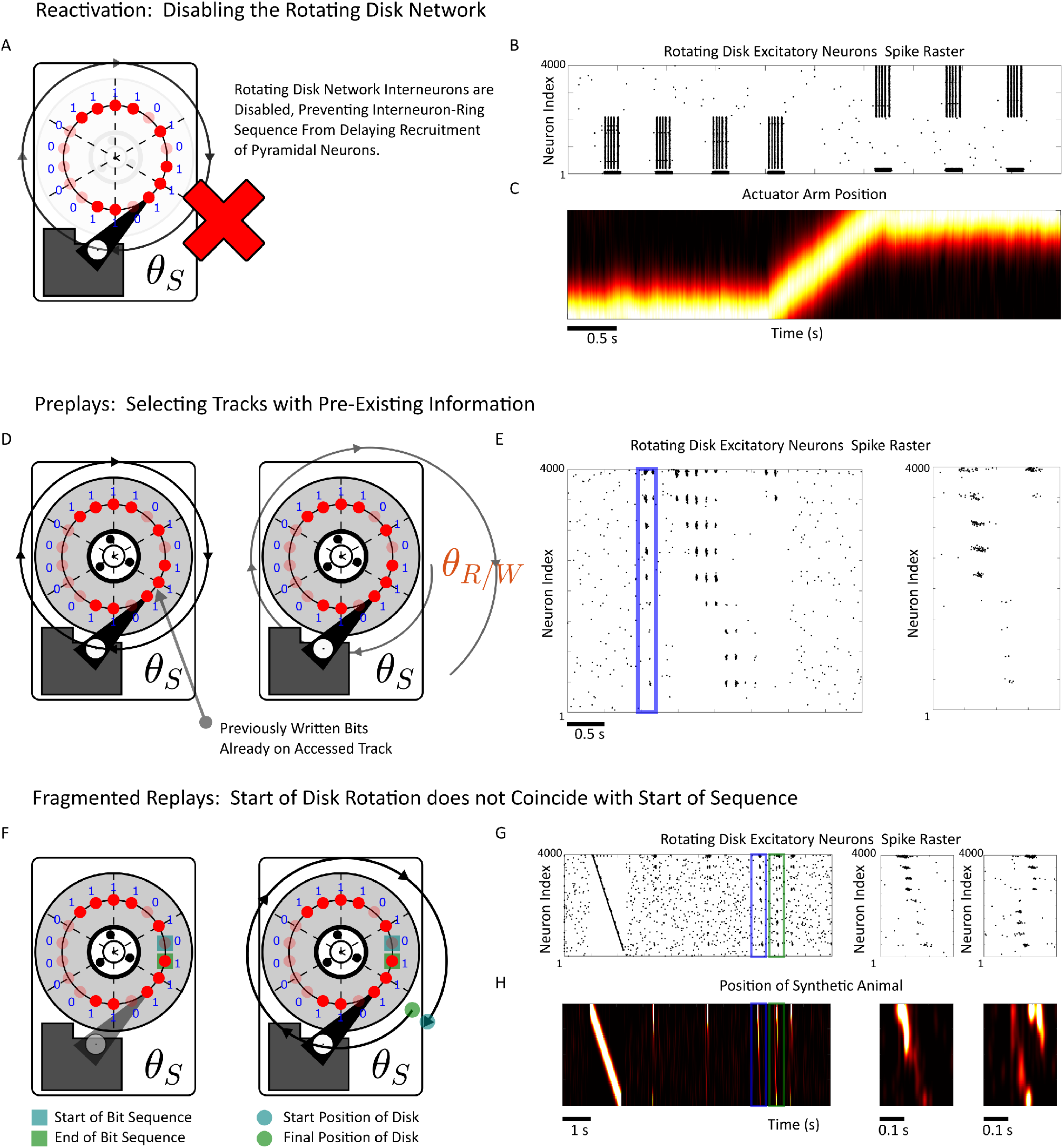
The Zoo of Hippocampal Data Access Methods **(A)** By disabling the rotating disk network interneurons, the spikes in the pyramidal neurons in the rotating disk network lose sequential content. **(B)** The simulated RD/AA network from Supplementary Figure 5A with the RD interneurons receiving a strong hyperpolarizing current to stop their recruitment. The pyramidal neurons now fire synchronized bursts in an event closer to reactivation rather than replay. **(C)** The position of the actuator arm, which can serve to bias which assembly becomes reactivated. **(D)** If a track already contains pre-written information, then new information can no longer be written to this track without the pre-existing information being deleted. Pre-plays may correspond to this scenario where tracks (SWRs) contain pre-existing information and are accessed during animal navigation. **(E)** In the NDD model, when information is already stored on a track/SWR, a compressed sequence (blue) occurs during a sharp-wave prior to sequential theta sequences during navigation. **(F)** For every sequence written to a track on a disk drive, there is a sequence start and end bit. If the disk rotation does not start with the sequence start bit, the sequence is accessed in a fragmented order. **(G)** Simulation of the NDD model with two separate replay events highlighted in blue and green. The first sequence, blue, corresponds to activating a track when the sequence starts. The second sequence, in green, corresponds to a mismatch between when the learned sequence starts and with which initial phase the rotating disk network is activated. **(H)** The decoded position of a synthetically generated animal. Replays are also decoded by a simple linear decoder, with a normal replay on the left (blue), and a fragmented replay on the right (green).

**Supplementary Figure 7:**
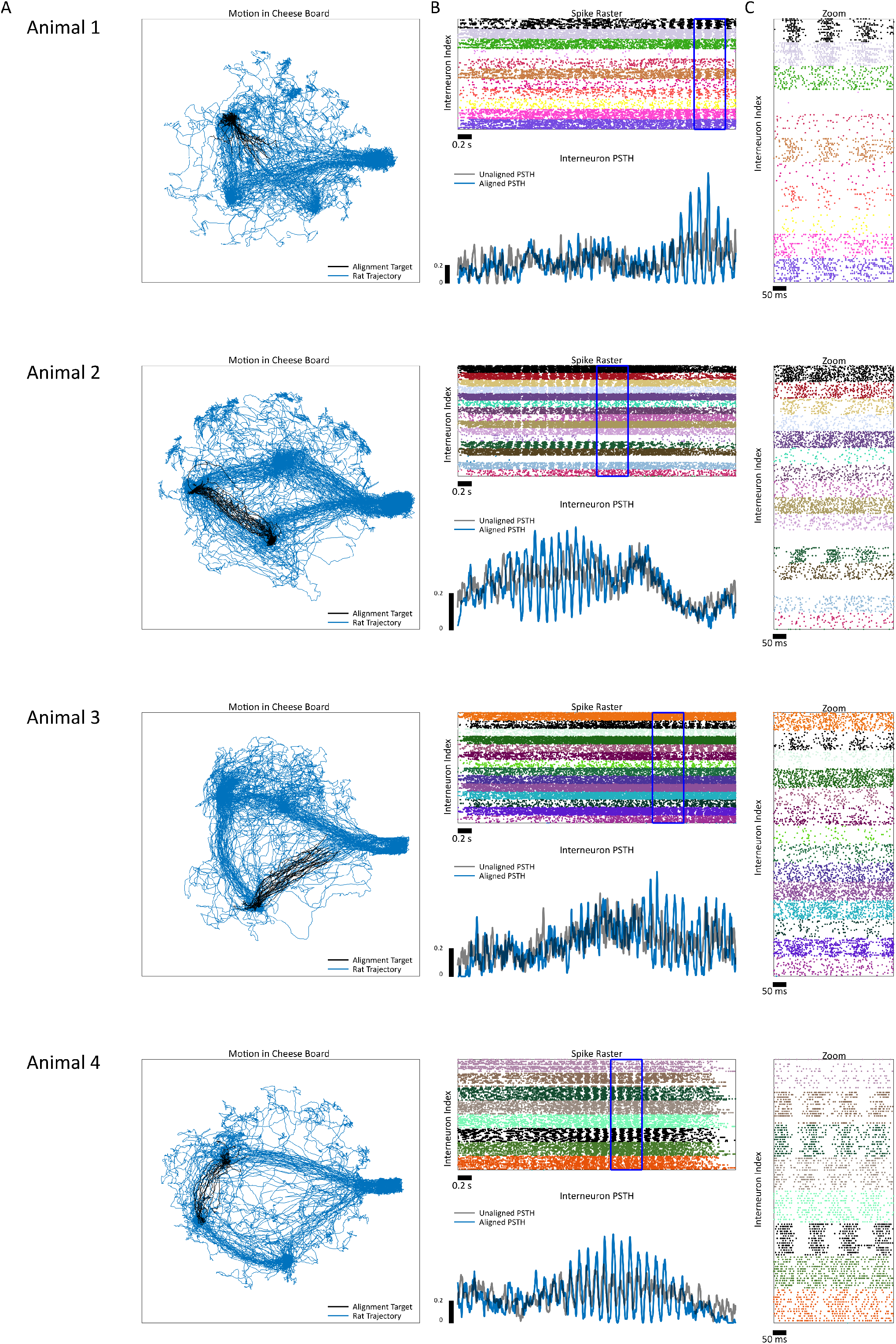
Time-Shift Alignment of Interneurons for 4 Different Animals **(A)** Motion trajectories used for each of the four animals as targets for maximum likelihood estimation based alignment of the spikes. **(B)** The interneurons in the recorded animal. The black interneuron was used as the target for alignment, with its determined time-shifts applied to all other interneurons. The kernel density estimate of the aligned (blue) and un-aligned (black) spike density for all interneurons is show. **(C)** A zoom of the interneurons during periods of high theta-locking in the spike-density. The zoomed segment corresponds to the blue box in (B).

**Supplementary Figure 8:**
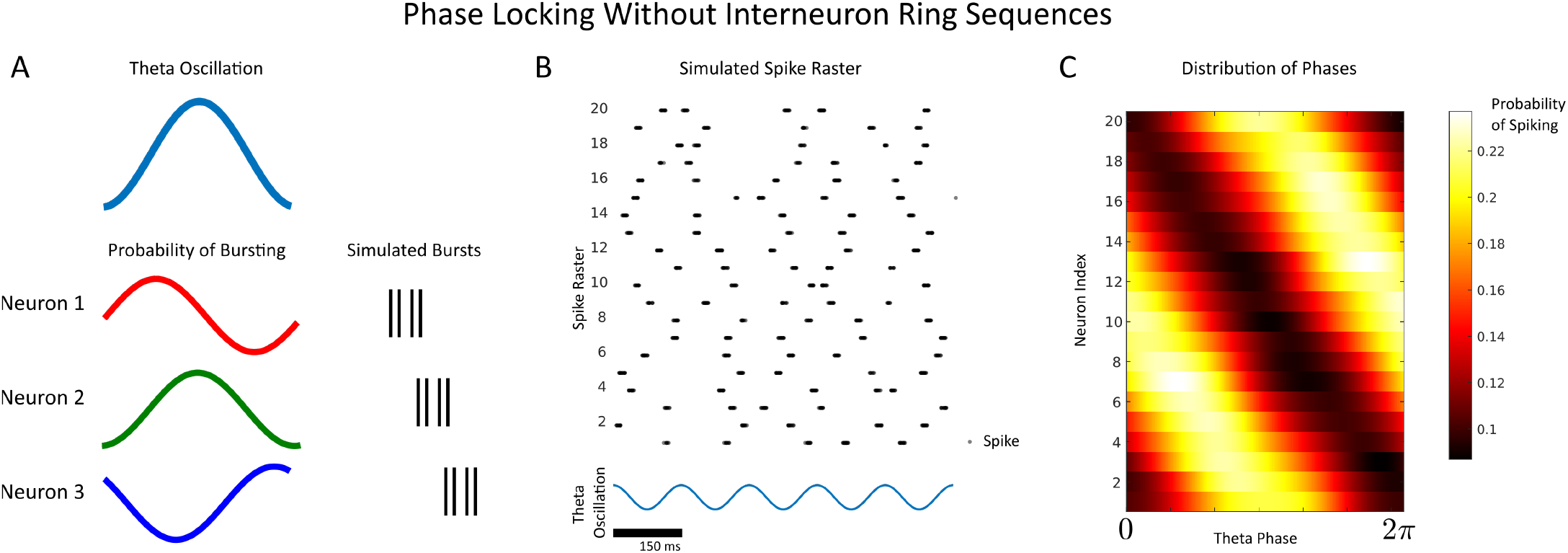
Preferred-Phase Firing without Interneuron Ring Sequences **(A)** In a theta-modulated poisson-spiking model, each neuron fires a burst of spikes with a theta modulated probability. The theta oscillation is used as a clock to force every cell to fire a burst, with the bursts elicited with a specific phase preference. **(B)** Despite phase-preferential firing of the neurons in this simulated system, a ring of bursts is not elicited. **(C)** Despite the lack of a ring of bursting, each neuron in the network exhibits phase locking of their spikes onto the theta oscillation. The phase locking forms a continuous ring, without interneuron-ring-sequences being elicited.

